# Guardian ubiquitin E3 ligases target cancer-associated APOBEC3 deaminases for degradation to promote human genome integrity

**DOI:** 10.1101/2024.04.23.590688

**Authors:** Irene Schwartz, Valentina Budroni, Mathilde Meyenberg, Harald Hornegger, Kathrin Hacker, Siegfried Schwartz, Zuzana Hodakova, Daniel B. Grabarczyk, Julian F. Ehrmann, Sara Scinicariello, David Haselbach, Jörg Menche, Tim Clausen, G. Elif Karagöz, Gijs A. Versteeg

**Author notes:** These authors contributed equally (Irene Schwartz, Valentina Budroni).

## Abstract

**Highlights:**

- RNA-free APOBEC3 (A3) can enter the nucleus, leading to genomic mutations.
- Three E3 ligases specifically bind the RNA-binding domain of nuclear A3s.
- Cancer-associated A3B and A3H-I are thereby targeted for proteasomal degradation.
- These E3 ligases thus act as genome guardians by limiting A3-mediated mutagenesis.

APOBEC family members play crucial roles in antiviral restriction. However, certain APOBEC3 (A3) proteins drive harmful hypermutation in humans, contributing to cancer. The cancer-associated A3 proteins are capable of transiting from the cytosol to the nucleus, where they can cause genome mutations. Here, we uncover a specific set of cellular pathways that protect genomic DNA from the major cancer-associated A3 proteins. Through genetic and proteomic screening we identify UBR4, UBR5, and HUWE1 as key ubiquitin E3 ligases marking cancer-associated A3B and A3H-I for degradation, thereby limiting A3-driven hypermutation. Mechanistically, UBR5 and HUWE1 recognize unoccupied A3 RNA-binding domains, thus promoting proteasomal degradation of APOBEC3 protein that is not engaged in its antiviral cellular function. Depletion or mutation of the E3 ligases in cells and human cancer samples increases A3-driven genome mutagenesis. Our findings reveal that UBR4, UBR5, and HUWE1 are crucial factors in a ubiquitination cascade that maintains human genome stability.

## Introduction

The human genome encodes seven members of the APOBEC3 (apolipoprotein B mRNA editing enzyme, catalytic polypeptide-like 3) family of cytidine deaminases^1^. These deaminases play a critical role in innate immunity against retro/lentiviruses by causing hypermutation of the viral cDNA^2,3^. The APOBEC3 (A3) family is under strong selective pressure in humans and other primates, with all seven members (A-H) possessing DNA C-to-U deaminase activity^4–7^. A3H is the oldest and most evolutionarily distant member of the A3 family and contains a unique zinc-coordinating motif in its deaminase domain^8,9^. In addition, it has the most haplotypes in the human population^1,10–12^. These haplotypes differ significantly in terms of their stability, subcellular localization and antiviral activity^13,14^.

In contrast to this host-beneficial function, previous studies have shown that several A3 family members can have detrimental effects in humans by driving hypermutation of cellular DNA. Such hypermutation has been documented in diverse cancer types, thereby contributing to a broader, disadvantageous mutational landscape within these tumors^12,15–18^.

Elevated levels of A3A, A3B, and A3H-I have been associated with mutagenesis in a range of cancers^15–17,19,20^. A3-mediated mutagenesis has been shown to drive some of the most prevalent mutational signatures in cancer, characterized by C-to-T transitions and clustered mutations (kataegis) at TCN trinucleotides^17,21–28^. APOBEC-associated mutational signatures have been identified in more than 70% of cancer types and around 50% of all cancer genomes^17,29,30^. These signatures are prominent in breast, lung, and bladder cancer, as well as other cancers^17,18,31–33^. A3A is overexpressed in a wide spectrum of human cancers and can induce kataegis and omikli, a form of extreme kataegis with more than 100 mutations per megabase^15–17,28,33^. A3B is overexpressed in many cancers and can generate APOBEC-specific SBS2 and SBS13 mutational signatures^17,34,35^. A3H occurs as several haplotypes in the human population, of which only the nuclear haplotype I (A3H-I) is associated with APOBEC signatures in breast and lung cancer, whereas the cytosolic haplotype II (A3H-II) is not^18,32^.

Although these A3 deaminases mutagenize ssDNA substrates, their localization and activity are controlled through binding to double-stranded secondary structures in cellular RNAs^36,37^. In infected cells, cytosolic A3 binding to secondary structures in the viral RNA genome is essential for packaging into progeny virions, and subsequent mutagenesis of the viral cDNA during reverse transcription^12–14,38,39^. A3F, A3G, and A3H-II have been reported to have strong virus restrictive properties^10–13^. Importantly, these A3 proteins are not turned-over by the proteasome, and as a consequence of their stability, accumulate at high steady-state intracellular protein concentrations^10–14,38^. In contrast, other A3 members which have been associated with hypermutation signatures in various cancers, such as A3A, A3B, and A3H-I, are predominantly nuclear. These A3s are rapidly turned-over by the proteasome, and consequently are present at low intracellular protein concentrations^15–17,19,20^. A3H-I instability is determined by a single nucleotide polymorphism (SNP) (R105G) that is associated with increased nuclear localization^18^. It remains unclear why this SNP results in increased nuclear localization and instability. Nuclear localization of A3A, A3B, and A3H-I has been proposed to contribute to cancer hypermutation as it promotes access to genomic DNA^18^.

The unstable A3 family members, though usually present at low nuclear levels, play a greater role in generating cancer-linked APOBEC mutational signatures than the more stable cytosolic variants^18,28^. However, it is unclear how the former group of A3s is more closely associated with cancer given their instability. We hypothesized that: **i)** deregulation of the limited protein concentrations of the unstable nuclear A3 members is likely sufficient to drive mutagenesis in cancers, **ii)** there are unidentified cellular factors that, under physiological conditions, maintain low nuclear A3 protein levels through active degradation, and **iii)** these cellular factors protect against cell-intrinsic genome mutagenesis by specifically keeping the cellular concentrations of potentially harmful nuclear A3 variants low. In addition, we predicted that the absence of these unidentified ‘guardian’ factors would unleash hypermutation through increased nuclear A3 protein levels that compromise host genome integrity.

Through genetic screening and proximity proteomics, we identified UBR4, UBR5, and HUWE1 as E3 ligases that ubiquitinate A3B and A3H-I, thereby targeting them for proteasomal degradation. Consistent with their genome-guardian roles, ablation or mutation of these E3 ligases in cancer cell lines and human cancer samples led to increased APOBEC3-driven hypermutation.

## Results

### Proteasomal degradation controls protein levels of cancer-associated A3s

Since the cancer-associated A3 family members, A3A, A3B, and A3H-I, only accumulate to low steady-state levels, we hypothesized that the cellular concentrations of these factors are controlled by protein degradation. To test this, constructs encoding all human A3 family members were delivered and expressed at similar steady-state protein levels (Fig. 1a). Subsequently, these cells were treated with the proteasome inhibitor epoxomicin (EPOX), and the effect on the various A3 proteins was determined by Western Blot (WB) analysis (Fig. 1a and 1b). Inhibition of proteasomal degradation significantly increased protein concentrations of the nuclear and cancer-associated A3A, A3B, and A3H-I proteins, indicating that their intracellular protein levels are substantially determined by proteasomal degradation. In contrast, protein levels of cytoplasmic A3s (A3D, A3F, A3G, and A3H-II), which are important for the innate immune response against retro/lentiviruses, were unaffected (Fig. 1a and 1b).

**Figure 1.**
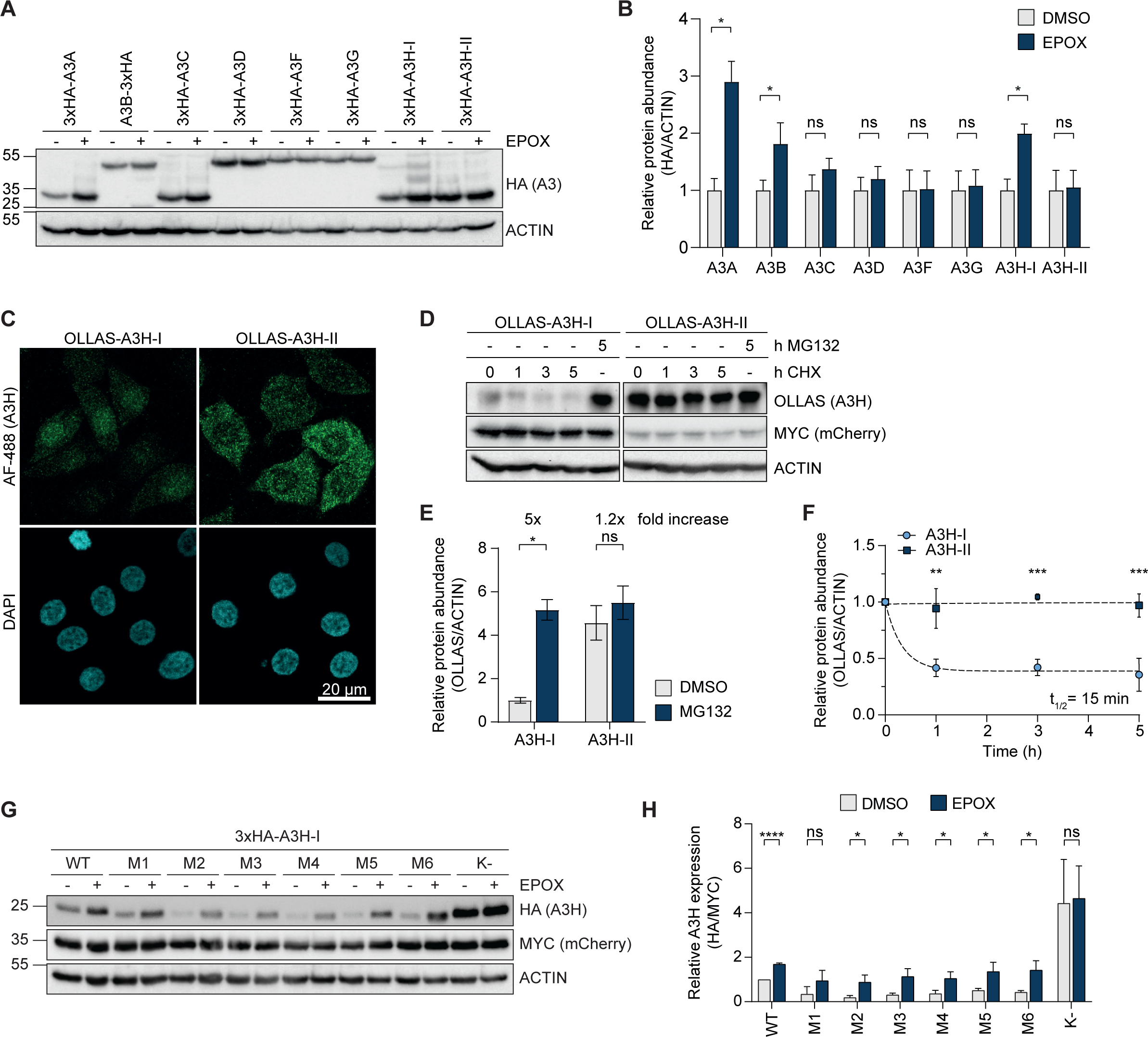
Proteasomal degradation controls protein levels of cancer-associated A3s. **(a)** HEK-293T cells were transfected with different amounts of 3xHA-tagged A3 expressing plasmids to achieve similar steady-state A3 protein levels. 24 h. post transfection, cells were treated for 16 h. with EPOX, and protein levels were analyzed by WB, and **(b)** quantified (means and SD, multiple unpaired t-tests, * p < 0.05, n = 5). **(c)** RKO cells stably expressing OLLAS-A3H-I/II were fixed, and their subcellular localization determined by immunofluorescence confocal microscopy; scale bar: 20 µm. **(d-f)** Lentiviral expression constructs encoding A3H-I or A3H-II were delivered to RKO cells at different integration rates to obtain comparable A3H-I and A3H-II protein levels in the presence of proteasome inhibitor. Polyclonal cell pools were treated with CHX or MG132 for the indicated times, **(d)** protein levels analyzed by WB, **(e)** relative A3H-I and A3H-II protein levels quantified by densitometry (means and SD, multiple unpaired t-tests, * p < 0.05, n = 2), and **(f)** single-step exponential decay curves were calculated, from which protein half-life was derived (means and SD, 2-way ANOVA, *** p < 0.0005, ** p < 0.005, n = 2). **(g-h)** HEK-293T cells were transfected with equal amounts of plasmids encoding the indicated MYC-mCherry-P2A-3xHA-tagged A3H-I mutants, in which multiple lysine residues were mutated to arginine. 36 h. post transfection, cells were treated with EPOX for 5 h., **(g)** protein levels determined by WB, and **(h)** quantified (means and SD, multiple unpaired t-tests, **** p < 0.0001, * p < 0.05, n = 3).

To further study the cellular mechanisms governing proteasomal degradation of nuclear, cancer-associated A3 protein levels in cells, we decided to use A3H as a model. The two predominant human A3H haplotypes represent each of the two identified phenotypes: A3H-I is nuclear, cancer-associated, and turned-over by the proteasome, whereas A3H-II is cytoplasmic and stable. Stability differences are reflected in much lower accumulation of A3H-I steady-state protein levels, as described for the endogenous protein in primary T lymphocytes^40^. This difference in stability is phenocopied upon exogenous expression in a wide variety of cell lines^12,40,41^. In line with previous reports^13,14,42^, some exogenous A3H-I localized to the cytosol but a substantial portion was nuclear, whereas A3H-II was mostly localized in the cytoplasm (Fig. 1c).

Subsequently, we set out to measure the protein stability of A3H-I and A3H-II. To this end, we generated polyclonal RKO (human colon carcinoma) and HeLa (human cervical adenocarcinoma) cell lines expressing a stable myc-tagged mCherry internal control, and either A3H-I or A3H-II through a P2A ribosomal skip site (Fig. S1a). To compensate for the differences in steady-state protein levels of the two A3H haplotypes, A3H-I cells were transduced with a higher virus-like particle concentration, and, therefore, express more of the mCherry control (Fig. 1d). A3H protein stability was then determined in a chase experiment with the translation inhibitor cycloheximide (CHX), or in the presence of proteasome inhibitor MG132 (Fig. 1d and S1b). Consistent with rapid proteasome-mediated turnover, A3H-I protein accumulated at much lower steady-state levels than A3H-II, and was rapidly depleted in the presence of cycloheximide (Fig. 1d and S1b, compare lanes 1 and 6). These levels were increased 4-to 5-fold upon proteasome inhibition (Fig. 1e and S1c). Confirming their different stabilities, A3H-I was degraded with a half-life of 15 min. in RKO cells (Fig. 1f), and 10 min. in HeLa cells (Fig. S1d), whereas A3H-II remained stable during the 5 h. chase period.

A3H-I degradation was exclusively dependent on proteasomal degradation, as steady-state protein levels of the two A3H haplotypes were unaffected by inhibitors of autophagy/lysosomal degradation (Fig. S1e). Likewise, no differences in mRNA stability in the presence of Actinomycin D (ActD) (Fig. S1f), nor secretion (Fig. S1g) were measured between the two A3H haplotypes, indicating that A3H-I protein levels were predominantly regulated through proteasomal degradation.

Consistent with this result, A3H-I and A3B (Fig. 1a-b) were ubiquitinated (Fig. S1h-i). Interestingly, while A3A was highly unstable (Fig. 1a-b), we consistently found it to be minimally ubiquitinated (Fig. S1h-i), which suggested that it may be regulated through a different - possibly ubiquitin-independent - mechanism than A3B and A3H-I.

To identify the residues important for A3H-I turnover, multiple lysine (K) residues in A3H-I were systematically grouped and mutated to arginine (R), based on their position in the A3H structure (Fig. S1j-k). All of these clustered K-to-R mutants accumulated at low steady-state protein levels, and were stabilized by proteasome inhibition, indicating that multiple lysines in different structural regions of A3H-I are likely important for its ubiquitination and degradation (Fig. 1g and 1h, Fig. S1a). In agreement with this conclusion and published data^43^, when all lysine residues were mutated to arginine, A3H-I accumulated at 4-fold higher steady-state protein concentrations and were no longer affected by proteasome inhibition (Fig. 1g-h). Together, these data indicate that protein levels of the nuclear, cancer-associated A3s, A3A, A3B, and A3H-I are regulated through proteasomal degradation, and that degradation of the model protein A3H-I depends on ubiquitination of multiple lysine residues.

### The E3 ligases UBR4, UBR5, and HUWE1 independently mediate turnover of A3B and A3H-I

The results described above positioned A3H-I as an excellent model to identify the cellular machinery that degrades it (and possibly other cancer-associated A3s), allowing its naturally occurring non-cancer-associated variant-A3H-II-to be used as a stable control to determine specificity.

To identify specific protein stability regulators of A3H-I, but not A3H-II, we set up a CRISPR-based genetic screening platform^44–46^. First, an RKO cell line was established harboring a doxycycline (DOX)-inducible Cas9-P2A-BFP construct, and an mCherry-A3H-II-P2A-GFP-A3H-I dual reporter (Dual-A3H-reporter), driven from an exogenous promoter (Fig. 2a). Inhibitor treatments confirmed that the two fluorophore-tagged A3H fusions phenocopied the protein stability pattern of their untagged counterparts (Fig. S2a).

**Figure 2.**
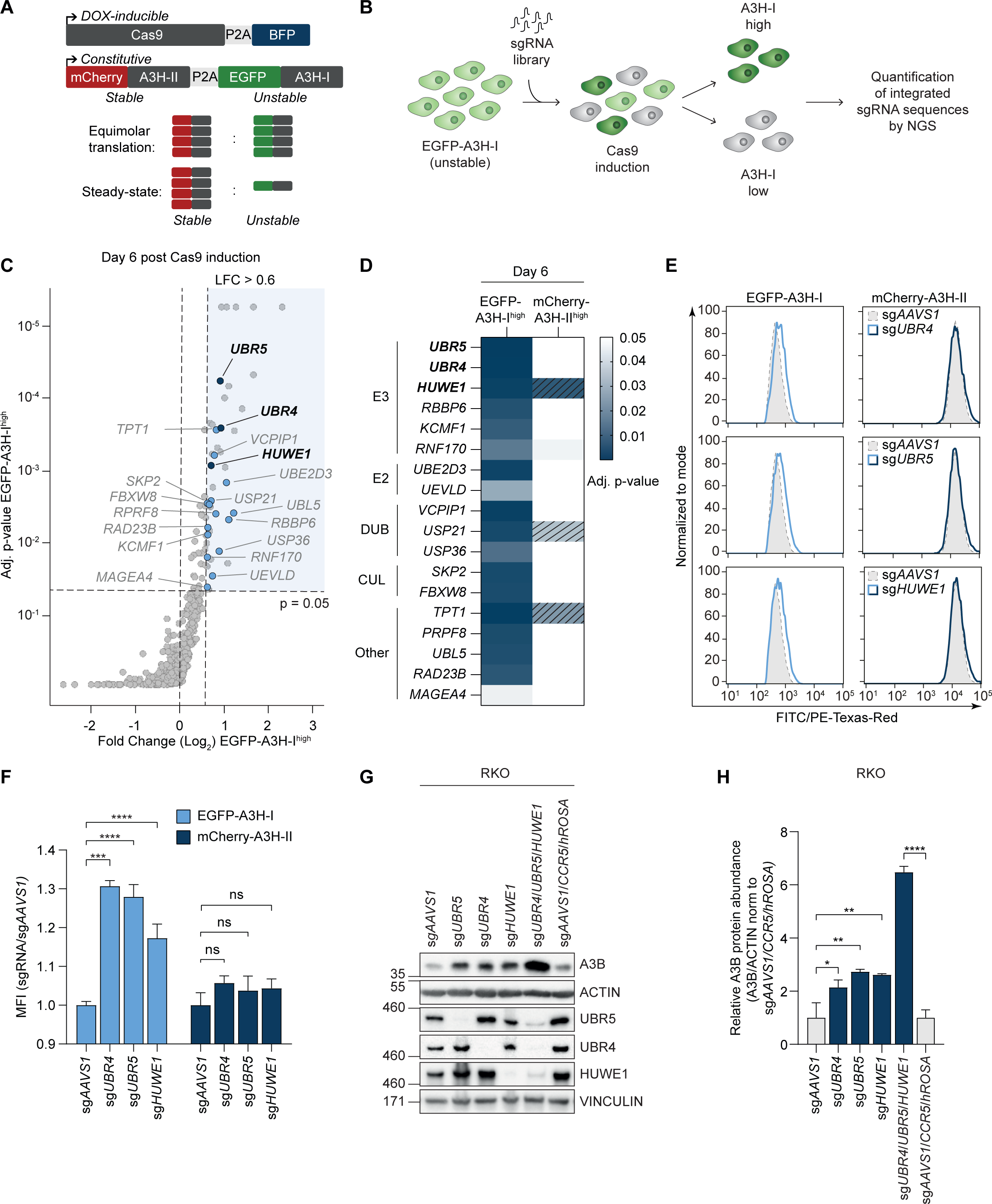
The E3 ligases UBR4, UBR5, and HUWE1 independently mediate turnover of A3B and A3H-I. **(a)** A monoclonal RKO cell line harboring a DOX-inducible Cas9-P2A-BFP expression construct and a constitutively expressed mCherry-A3H-II-P2A-EGFP-A3H-I dual-reporter (RKO-DOX-Cas9-dualA3H) was established. Unstable EGFP-A3H-I and stable-mCherry-A3H-II are synthesized in equimolar amounts, yet EGFP-A3H-I accumulates at low steady-state levels due to proteasomal degradation. **(b)** Schematic representation of CRISPR/Cas9 FACS-based screening procedure. RKO-DOX-Cas9-dualA3H cells were transduced with an sgRNA library targeted at ubiquitin-proteasome- and autophagy-linked factors. After selection of sgRNA-positive cells with G418, Cas9 expression was induced with DOX for 3 and 6 days, followed by sorting of cells with the 1-2% highest and lowest EGFP or mCherry fluorescence by flow cytometry. Integrated sgRNA sequences were identified by next generation sequencing (NGS), and their differential abundance relative to non-sorted cells analyzed. **(c)** Targeted genes enriched in EGFP-A3H-I^high^ sorted cell populations 6 days post Cas9 induction. **(d)** Heatmap of top genes on log_2_ fold-change and p-value grouped by functional categories. Genes enriched in EGFP-A3H-I^high^ cell populations 6 days post Cas9 induction with a log_2_ fold-change >0.6 which were not enriched in mCherry^high^ or GFP^low^ on either day 3 (LFC > 0.45) or day 6 (LFC > 0.6). Adjusted p-values are based on MaGECK analysis of three independent replicate sorts. Dashed lines indicate a log_2_ fold-change < 0.6. E3 (E3 ligases), E2 (E2 conjugating enzymes), DUB (Deubiquitinases), CUL (components of Cullin-RING ubiquitin ligase complex). **(e)** Monoclonal RKO-Dox-Cas9-dualA3H cells were transduced with individual sgRNAs targeting the indicated genes. Subsequently, Cas9 expression was induced by DOX for 6 days and EGFP-A3H-I and mCherry-A3H-II mean fluorescence intensity (MFI) analyzed by flow cytometry, and **(f)** quantified (means and SD, one-way ANOVA, **** p < 0.0001, *** p < 0.0005, n = 3). **(g)** RKO cells harboring DOX-inducible Cas9 were transduced with sgRNAs targeting *UBR4*, *UBR5*, or *HUWE1,* individually or in combination, and sorted for sgRNA-positive cells. Gene editing was induced with DOX for 6 days, after which endogenous A3B protein levels were determined by WB, and **(h)** quantified (means and SD, one-way ANOVA, **** p < 0.0001, ** p < 0.005, * p < 0.05, n = 2).

To screen for regulators of A3H-I stability, the Dual-A3H-reporter cell line was transduced with a ubiquitin-focused lentiviral sgRNA library, Cas9 expression was induced, and cells with the highest or lowest 1-2% EGFP or mCherry fluorescence (EGFP^high^, mCherry^high^, EGFP^low^, and mCherry^low^) were collected after 3 and 6 days. Genomic DNA was isolated from the collected cells, and integrated sgRNA-coding sequences quantified by NGS (Fig. 2b).

Several sgRNAs were specifically enriched in EGFP-A3H-I^high^ sorted cells, which did not score in the mCherry-A3H-II^high^ sorted pool (Fig. 2c-d). Among the top hits were sgRNAs targeting several E3 ligases, E2 conjugating enzymes, DUBs/proteases and other degradation-associated genes. The top E3 ligases scored on day 6 were *UBR4*, *UBR5*, and *HUWE1. HUWE1* was also one of the top hits on day 3 (Fig. S2b-c). Interestingly, we also found the gene encoding the E2 conjugating enzyme UBE2D3, an E2 conjugase known to work with UBR5 and HUWE1, as one of the top hits for both days (Fig. 2c-d and S2c-d). These results identified UBR4, UBR5, and HUWE1 as strong, specific candidates mediating A3H-I degradation.

To test the validity and specificity of the screen results, each of these E3 ligases was targeted in isolation. Individual ablation of each of the E3 ligases significantly increased EGFP-A3H-I levels in the Dual-A3H-reporter screening cell line (Fig. 2e-f), as well as a polyclonal cell pool expressing the same construct (Fig. S2d), whereas mCherry-A3H-II fluorescence was not affected. We further tested whether the increased A3H-I steady-state protein levels in the E3 ligase knock-out cells stemmed from increased protein stability, by inhibiting translation with CHX. Therefore, we generated RKO-DOX-Cas9-MYC-mCherry-P2A-3xHA-A3H-I cells harboring a triple-E3 ligase knock-out, treated them with CHX, and monitored A3H-I protein levels over time by WB (Fig. S2e-f). Ablation of all three E3 ligases increased A3H-I half-life from 21 min. to 1.9 h. (Fig. S2f), indicating that A3H-I degradation is compromised in the absence of UBR4/UBR5/HUWE1.

Since we found that A3B is also turned-over in a proteasome-dependent manner (Fig. 1a-b), we reasoned that the same cellular machinery may be employed to keep its steady-state levels low. Therefore, we set out to determine whether *UBR4*, *UBR5*, or *HUWE1* ablation would also influence the abundance of A3B. Individual knock-out of *UBR4*, *UBR5,* and *HUWE1*, increased endogenous A3B protein levels by 2.5 – 4-fold in RKO cells (Fig. 2g-h) as well as in THP-1 monocytic cells (Fig. S2g-h), without affecting its mRNA levels (Fig. S2i). Consistent with the hypothesis that the E3 ligases specifically degrade nuclear A3 members, the predominantly cytoplasmic endogenous A3G remained unaffected by the E3 knock-outs in THP-1 cells (Fig. S2g-h). In RKO cells, stability of endogenous A3G could not be evaluated due to low A3G expression in these cells.

Importantly, knock-out of each of the individual ligases increased endogenous A3B protein levels by 2-fold in each case (Fig. 2g-h), whereas simultaneous targeting of *UBR4*, *UBR5*, and *HUWE1*, strongly increased A3B levels in an additive fashion to 6-fold in comparison to its matching controls in which three safe harbor loci were targeted (Fig. 2g-h). From these non-epistatic genetic interactions, we concluded that UBR4, UBR5, and HUWE1 each target A3B independently of the other two E3 ligases.

Collectively, these results identified the E3 ligases UBR4, UBR5, and HUWE1 as important functional effectors of A3H-I degradation. This degradation-dependence is conserved for cancer-associated A3B and is relevant in different human cell types. Our results indicate that the three E3 ligases likely function redundantly, and thus poly-ubiquitinate A3B, and possibly also A3H-I, independently of each other.

### UBR5 and HUWE1 form a complex with A3H-I and other unstable A3 deaminases in cells

The above data argue that UBR4, UBR5, and HUWE1, identified through our specific genetic screen, can directly or indirectly mediate A3 degradation. To complement the genetic screen, we set out to also identify specific physical interaction partners of the unstable A3H-I. We reasoned that overlapping factors identified in both approaches would be strong candidates for factors that directly recognize A3B and A3H-I as ubiquitination substrates.

For unbiased identification of specific A3H-I interactors, TurboID (TID) proximity labeling proteomics was performed (Fig. 3a). Three different RKO cell lines stably expressing either DOX-inducible **a)** TID-A3H-I, **b)** TID-A3H-II, or **c)** TID-GFP as a control were generated. Similar expression of each of the three constructs was achieved by inducing transgene expression with different concentrations of DOX. Subsequently, proteins proximal to the TurboID fusion proteins were covalently labeled by addition of biotin, after which biotinylated proteins were isolated and identified by nLC-MS/MS.

**Figure 3.**
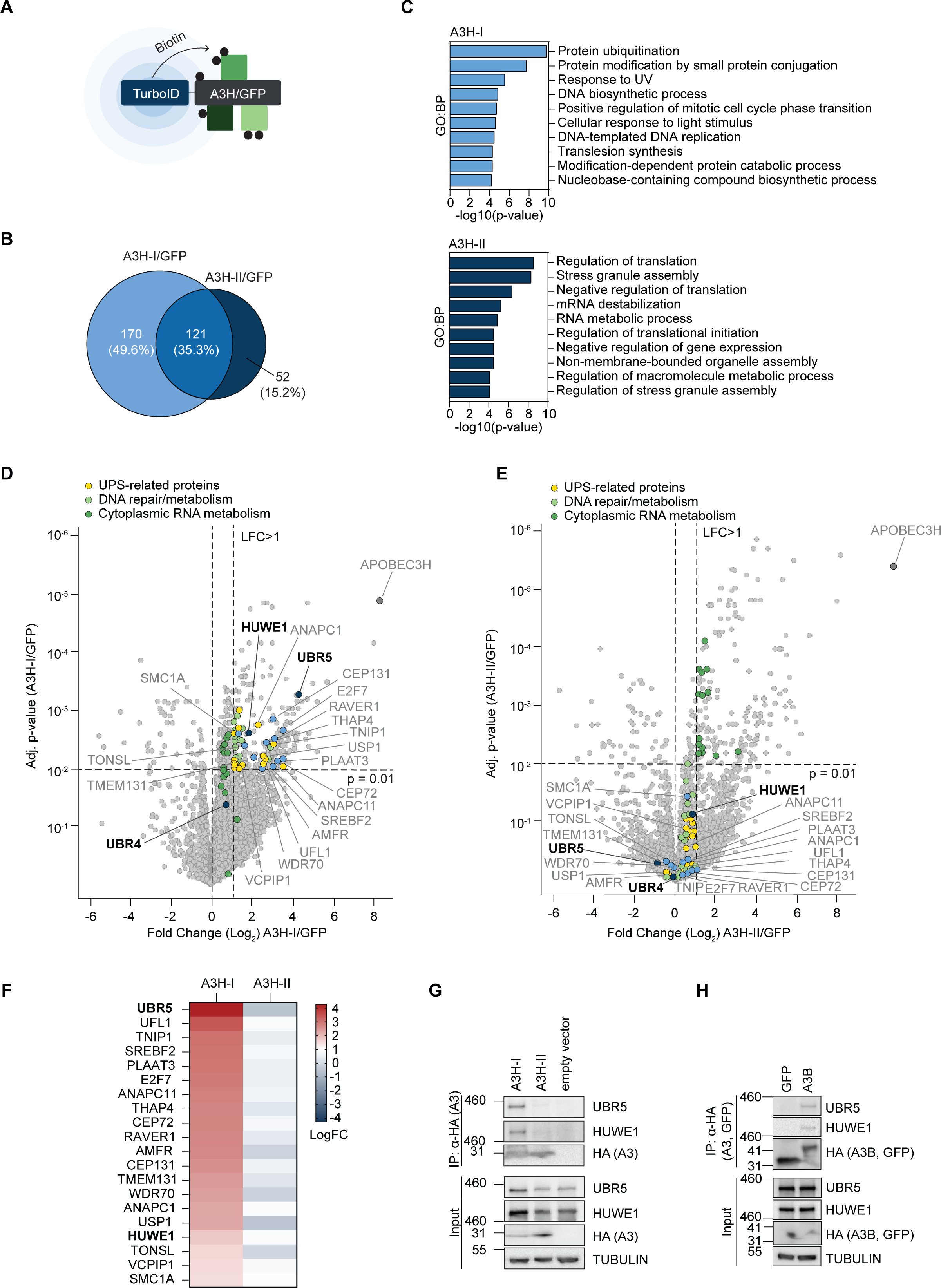
UBR5 and HUWE1 form a complex with A3H-I and other unstable A3 deaminases in cells. **(a)** Overview of TurboID proximity labeling principle. **(b)** Polyclonal RKO-DOX-TID-A3H-I/II/GFP were treated with different concentrations of DOX for 2 days to achieve similar protein levels. Subsequently, cells were treated with EPOX for 5 h. and the culture media supplemented with biotin during the last 15 min. Biotinylated proteins were purified under denaturing conditions and quantified by nLC-MS/MS. Differentially enriched proteins in A3H-I/GFP (light blue, LFC > 1, p-value < 0.01) and A3H-II/GFP (dark blue, LFC > 1, p-value < 0.01) were compared. **(c)** GO terms for biological processes (GO:BP) of differentially enriched proteins in A3H-I/GFP (light blue, LFC > 1, p-value < 0.01, input: 170 factors derived from (b)) and A3H-II/GFP (dark blue, LFC > 1, p-value < 0.01, input: 52 factors derived from (b)). **(d)** Differential expression of TID-A3H-I, or **(e)** TID-A3H-II interactors relative to TID-GFP (n = 3). Proteins specifically enriched for either A3H-I or A3H-II identified in (a) are indicated in color. **(f)** Heatmap representation of enriched proteins in TID-A3H-I samples, or TID-A3H-II samples relative to TID-GFP. Top 20 A3H-I-specific proteins displayed according to the following criteria: A3H-I/GFP LFC > 1, p-value < 0.01, which are not enriched in A3H-II/GFP LFC > 1, p-value < 0.01. **(g-h)** HEK-293T cells were transiently transfected with different amounts of plasmids encoding **(g)** 3xHA-A3H-I/II, or **(h)** 3xHA-GFP or A3B-3xHA to achieve similar steady-state protein levels. Subsequently, 3xHA-tagged proteins were immunoprecipitated, and their interaction with endogenous UBR5 and HUWE1 determined by WB.

We first compared the interactomes of A3H-I and A3H-II relative to the TID-GFP control. In line with a recent publication^47^, both A3H haplotypes share many interactors (Fig. 3b), mainly comprising Gene Ontology (GO) terms for biological processes representing regulation of translation, RNA metabolism, and RNA processing (Fig. S3a), which are amongst the top scoring hits in both A3H haplotypes.

However, analysis of A3H-I-specific interactors identified protein ubiquitination and DNA-metabolism related processes as the top ranked biological process GO terms (Fig. 3c). Processes related to A3H-II-specific included translation, stress granule assembly, and RNA metabolic processes (Fig. 3c), terms which are partially shared with the A3H-I interactome (Fig. S3a). This reinforces the above results, which indicate that only A3H-I is targeted by the ubiquitin-proteasome system. In addition, the enrichment for DNA-metabolism related processes indicates that A3H-I may have access to DNA. As expected, top-ranked GO terms for cellular compartments were enriched for nuclear terms in the case of the A3H-I-specific interactome. In contrast, A3H-II-linked terms predominantly included cytoplasmic stress granules and P-bodies (Fig. S3b), underlining the propensity of stable A3s to form high-molecular-weight RNP complexes^48–51^.

Consistent with a high degree of shared interaction partners, most interactors were not unique to either the A3H-I/GFP or A3H-II/GFP haplotype (Fig. 3d-e; unlabelled data points in upper-right quadrant). Amongst the specific proteins significantly enriched in the A3H-I samples were the E3 ligases UBR5 and HUWE1 (Fig. 3d-f), which were also identified as specific genetic interactors of A3H-I (Fig. 2c-d). Although UBR4 peptides were detected by nLC-MS/MS, they were not significantly enriched in any one sample. In line with these MS results, WB analysis detected UBR5 and HUWE1 in streptavidin-enriched lysates from TID-A3H-I expressing cells, but not in lysates from TID-A3H-II or TID-GFP controls (Fig. S3c). Consistent with the proximity labeling results, endogenous UBR5 and HUWE1 specifically interacted with exogenously expressed A3H-I (Fig. 3g) and A3B (Fig. 3h) but did not co-IP with the stable A3H-II haplotype (Fig. 3g).

Together, these data show that UBR5 and HUWE1 specifically interact with A3B and A3H-I in cells and may thus directly ubiquitinate them. In contrast, UBR4 was not identified as a specific A3B/A3H-I interactor in cells. UBR4 may either have an indirect effect on A3B and A3H-I stability, or its interaction with these A3s may be too transient to detect in cell-based assays.

### RNA binding protects A3 from E3 ligase binding and ubiquitination, thus promoting stability in cells

RNA binding plays an important regulatory role in A3 localization and antiviral activity^39,48,52,53^. The antiviral family members A3H-II and A3G are retained in the cytoplasm through RNA binding^39,48,52,53^. A3H haplotype I and II differ by three amino acids (Fig. S4a), with a single glycine at position 105 determining the difference in stability^14,18,32^. In line with these data, mutation of G105R in A3H-I turns it from unstable and partially nuclear, to stable and predominantly cytosolic, phenocopying A3H-II (Fig. S4b-d). Conversely, R105G mutation renders A3H-II unstable and partially nuclear, thereby phenocopying A3H-I (Fig. S4b-d). Previous work has suggested that the R105G mutation may render the A3H protein less capable of binding dsRNA structures, leading to diminished antiviral potential^14,18,32^. Furthermore, introducing the G105R into A3H-I, strongly reduced its ubiquitination (Fig. S4e) and interaction with UBR5 and HUWE1 (Fig. S4f). Conversely, the A3H-II R105G mutation resulted in its ubiquitination (Fig. S4e) and association with UBR5 and HUWE1 (Fig. S4f).

Based on these and prior observations^14,18,32^, we hypothesized that lack of RNA binding leads to the re-localization of cytoplasmic A3H-II to the nucleus and renders it a substrate for proteasomal degradation. Therefore, we tested previously characterized A3H-II mutants (W115A and R175/176E)^39^, which harbor mutations in the α6 helix and the structurally adjacent loop 1, which mediate RNA binding. In line with previous reports^39,48,52,53^, lack of RNA binding decreased cytosolic A3H-II levels and increased nuclear localization (Fig. S4g). In contrast to the stable wild-type (WT) A3H-II, the RNA-binding mutants were highly unstable: their levels decreased rapidly upon translation inhibition yet stabilized in the presence of a proteasome inhibitor (Fig. 4a-b). These results indicated that RNA binding affects two characteristics of cellular A3: **i)** it retains it in the cytosol, and **ii)** it prevents its degradation.

**Figure 4.**
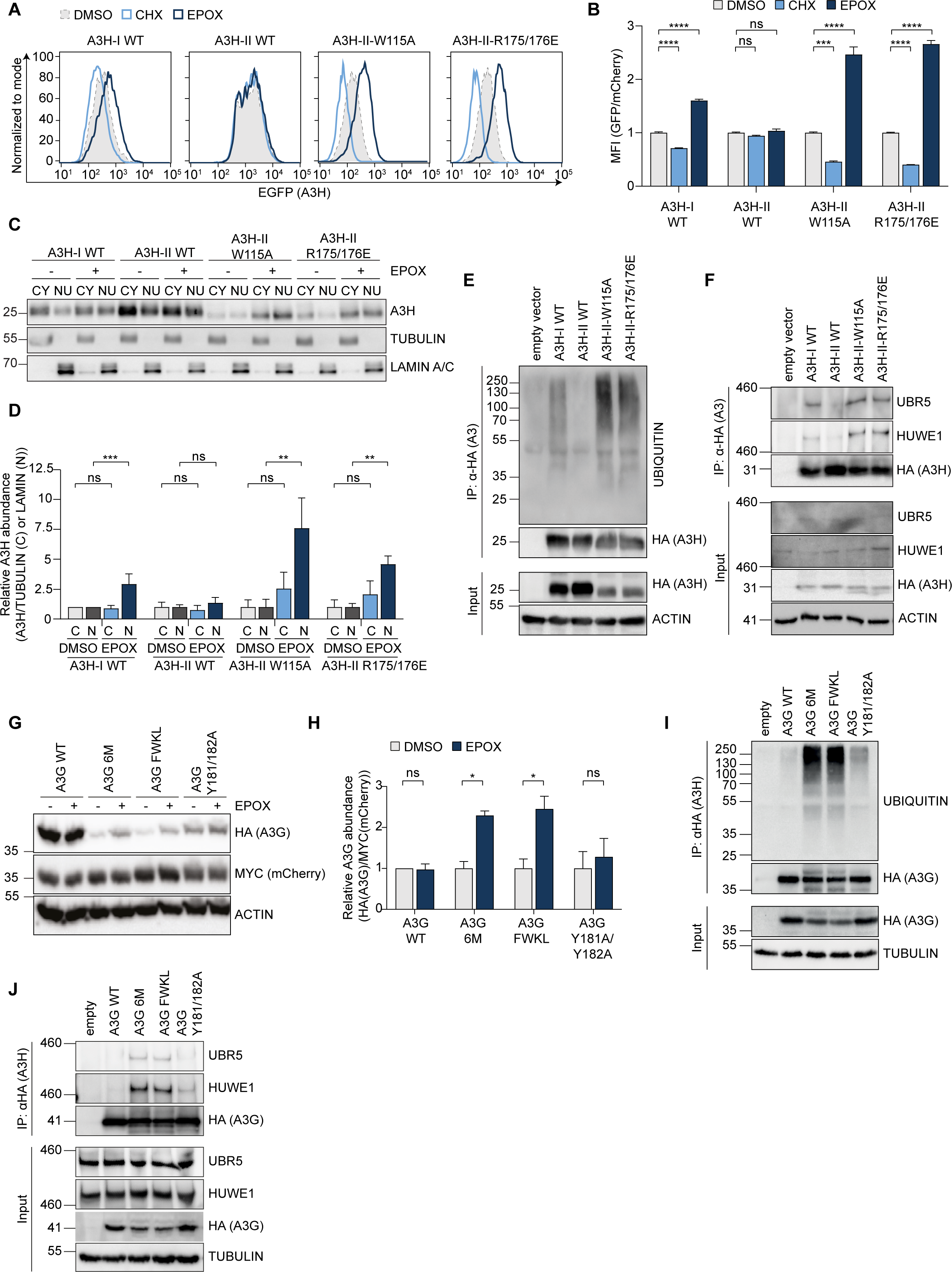
RNA binding protects A3 from E3 ligase binding and ubiquitination, thus promoting stability in cells. **(a)** RKO-mCherry-P2A-EGFP-A3H cells expressing the indicated EGFP-tagged A3H variants were treated for 5 h. with EPOX or CHX, after which mCherry and EGFP-A3H abundances were measured by flow cytometry, and **(b)** the MFI quantified (means and SD, one-way ANOVA, **** p < 0.0001, *** p < 0.0005, n = 3). **(c)** HEK-293T cells were transfected with different amounts of plasmids expressing the indicated 3xHA-tagged A3H constructs to achieve similar steady-state protein levels. Following 5 h. of EPOX treatment, whole cell extracts (Fig. S4h), cytoplasmic (C) and nuclear (N) fractions were extracted, analyzed by WB, and **(d)** quantified (means and SD, one-way ANOVA; *** p < 0.0005, ** p < 0.005; n = 3). **(e-f)** HEK-293T cells were transfected with different amounts of plasmids expressing the indicated 3xHA-tagged A3H constructs to achieve similar protein levels. Following 5 h. of EPOX treatment, 3xHA-tagged proteins were immunoprecipitated, and **(e)** their ubiquitination or **(f)** interaction with UBR5 and HUWE1 analyzed by WB. **(g)** mCherry-P2A-3xHA-tagged A3G WT and A3G RNA-binding mutants (6M (F126Y, W127S, K180A, I183A, L184A, I187A), FWKL (F126Y, W127S, K180S, L184S) and Y181A/Y182A) were transiently expressed in HEK-293T cells, followed by 5 h. of EPOX treatment and analysis of protein levels by WB, and **(h)** quantified. Multiple unpaired t-tests, * p < 0.05; n = 3). **(i-j)** HEK-293T cells were transfected as in (g), but with different amounts of plasmid constructs to achieve similar steady-state protein levels, followed by EPOX treatment for 5 h. and immunoprecipitation of 3xHA-tagged proteins from the cell lysates. Their **(i)** ubiquitination, or **(j)** interaction with UBR5 and HUWE1 was analyzed by WB.

Next, we asked in which compartment the respective A3H mutants are degraded. For this experiment, A3H WT and RNA-binding mutant constructs were expressed at similar levels, after which their accumulation upon proteasome inhibition was assessed in the nuclear and cytosolic cell fractions (Fig. 4c-d). As expected, A3H-II protein levels in whole cell extracts were not affected by proteasome inhibition, whereas it significantly increased the levels of the unstable A3H-I and A3H-II RNA-binding mutants (Fig. S4h-i). Analysis of fractionated samples showed that A3H-I and the RNA-binding mutants of A3H-II predominantly accumulated in the nucleus when their degradation was blocked (Fig. 4c-d), indicating that unstable A3H variants are degraded in the nucleus. Moreover, RNA-binding mutants of A3H-II were strongly ubiquitinated (Fig. 4e) and co-immunoprecipitated with UBR5 and HUWE1 (Fig. 4f), indicating that loss of RNA binding renders A3H-II a substrate for degradation by the identified E3 ligases. These data indicated that RNA binding may be a general mechanism by which A3 protein degradation is prevented, and may be the reason why other family members, such as A3G, are stable. To test this hypothesis, we analyzed the stability of WT A3G compared to three A3G RNA-binding mutants^54^ in cells. Consistent with comparable RNA-binding mechanisms across the A3 family, structural alignment with A3H showed that the helix region of A3G CD1-containing mutations superimposed with the α6 helix of A3H-II, identifying it as a likely determinant for RNA binding (Fig. S4j). In contrast to WT A3G, steady-state protein levels of A3G RNA-binding mutants increased upon proteasome inhibition, underlining the importance of the integrity of this helix, its ability to bind RNA, and formation of high molecular weight complexes for protein stability (Fig. 4g-h). Notably, A3G-Y181/182A was stabilized less upon EPOX treatment (Fig. 4g-h), possibly resulting from incomplete disruption of the α-helix by these mutations.

In agreement with the above data, which argue that RNA binding prevents recognition of A3 as substrates for degradation, the A3G 6M and FWKL mutants were ubiquitinated (Fig. 4i) and co-immunoprecipitated with UBR5 and HUWE1 (Fig. 4j), whereas their WT counterparts did not. Lower levels of A3G Y181/182A ubiquitination and E3 interaction (Fig. 4i-j) correlated with reduced instability of this mutant compared to WT protein (Fig. 4g-h), potentially stemming from differences in their RNA-binding.

Collectively, these results show that the RNA-binding ability of A3H-II and A3G are important for their stability. Absence of RNA binding enables UBR5 and HUWE1 engagement, ubiquitination, and eventual proteasomal degradation in cells.

### RNA binding by APOBEC3 proteins prevents their recognition and ubiquitination by E3 ligases *in vitro*

Cell-based assays showed that impaired RNA binding renders A3 proteins substrates for proteasomal degradation. Therefore, we hypothesized that the identified E3 ligases recognize the unoccupied RNA-binding surface of A3 proteins, and that RNA binding thus prevents ubiquitination. However, RNA binding has two confounding effects in cells as it influences both A3 localization, and stability. To test the hypothesis that the E3 ligases recognize unengaged RNA-binding interfaces, we therefore set out to analyze direct E3 ligase targeting of A3H in a cell-free *in vitro* reconstituted ubiquitination system.

First, A3H-I and A3H-II were purified from *E. coli* as previously described^39,52,55,56^. Despite several attempts during purification to remove co-purified bacterial RNA bound to A3H-I WT and A3H-II WT with high salt or extensive RNase A treatment, we were unable to purify these proteins without any residual bound RNA, as both proteins became insoluble and precipitated upon complete removal of RNA (not shown). For that reason, we additionally purified an A3H-II RNA-binding mutant (A3H-II-RBM) from *E. coli*, which has been previously reported to purify as an RNA-free monomer (Fig. 5a)^39^. In cells, this mutant phenocopied the stability of other A3H-II RNA-binding mutants (W115A and R175E/176E) (Fig. S5a-b).

**Figure 5.**
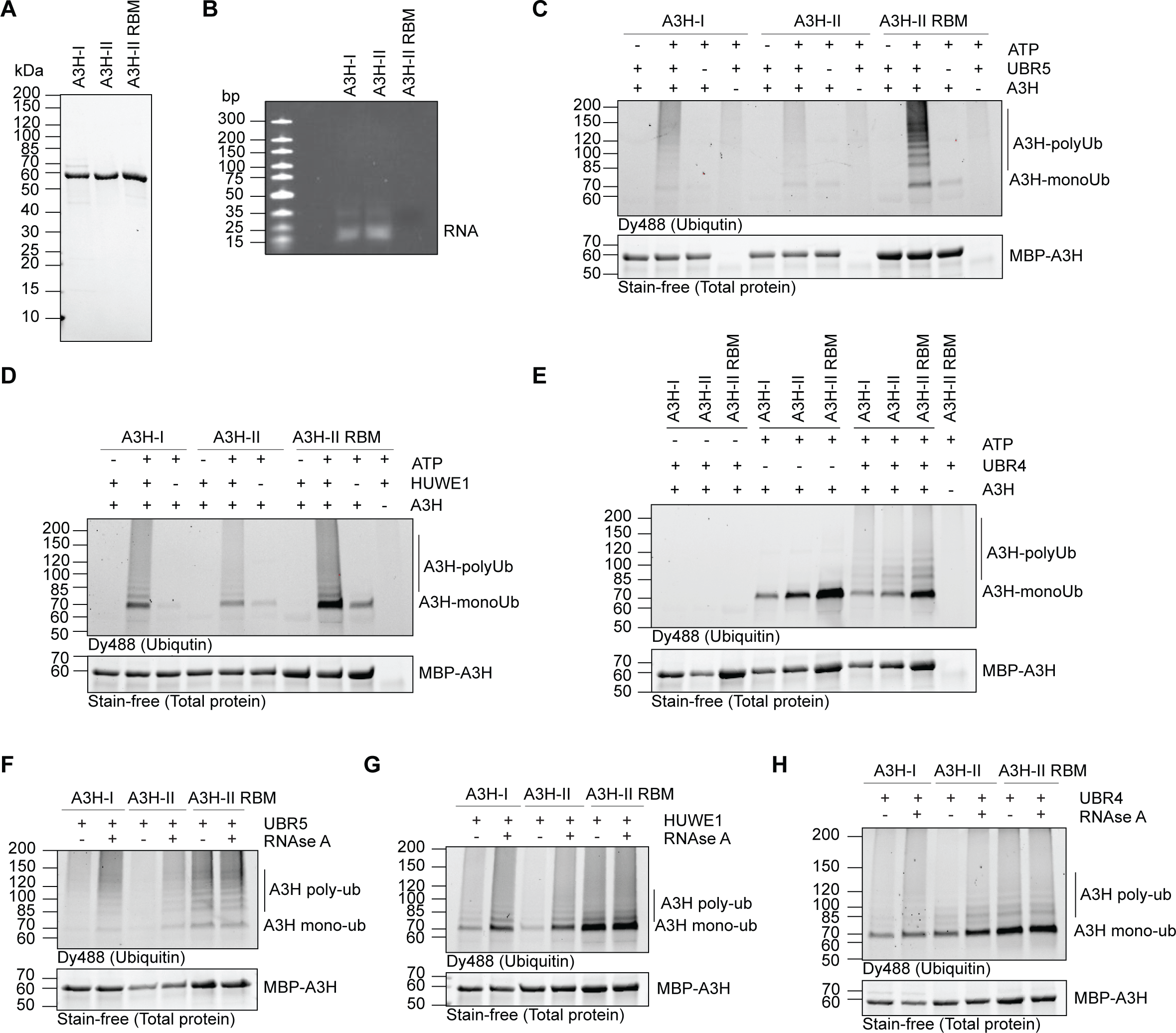
RNA binding by APOBEC3 proteins prevents their recognition and ubiquitination by E3 ligases *in vitro*. **(a)** 10x-His-MBP-A3H-I, 10x-His-MBP-A3H-II and 10xHis-MBP-A3H-II-RBM (E56A/ W115A/ R175E/ R176E) were expressed in *E. coli*, purified by HisTrap and subsequent gel filtration, after which purified proteins were analyzed by PAGE and Coomassie staining. **b)** Purified proteins were analyzed by denaturing Urea-TBE PAGE followed by SYBR Gold staining. **(c-e)** *In vitro* ubiquitination assays were performed with recombinant **(c)** UBR5, **(d)** HUWE1, or **(e)** UBR4 and WT A3H-I/II or A3H-RBM as substrates, and in the presence of DyLight488-labeled recombinant ubiquitin. Subsequently, A3H was immunoprecipitated using anti-MBP-coupled beads and the ubiquitination pattern visualized by fluorescent imaging for DyLight488. **f-h)** *In vitro* ubiquitination assay with recombinant **f)** UBR5, **g)** HUWE1, **h)** UBR4 and A3H-I/II WT or A3H-RBM in the absence or presence of RNase A. Ubiquitinated A3H was visualized as in (c-e).

A3H-I eluted exclusively as a monomer in size exclusion chromatography (SEC), whereas A3H-II eluted both as monomer and dimer (Fig. S5c). However, both proteins were bound to substantial amounts of short RNA fragments, as visualized by denaturing PAGE analysis (Fig. 5b), consistent with previous reports^39,52,55,56^. In contrast, A3H-II-RBM eluted as a monomer, free from any bound RNA (Fig. 5b and S5c).

Our cell-based data suggested that UBR5, HUWE1, and UBR4 poly-ubiquitinate A3H-I in cells, likely independently of one other (Fig. 2g-h). To validate these results in a cell-free system, we performed *in vitro* ubiquitination assays using purified proteins and either A3H-I, A3H-II or the RNA-free A3H-II-RBM mutant as substrates. Consistent with our cell-based data, UBR5 and HUWE1 independently poly-ubiquitinated A3H-II-RBM and A3H-I (Fig. 5c-d). Both enzymes polyubiquitinated the RNA-free A3H-II-RBM mutant most strongly, followed by RNA-bound A3H-I, with A3H-II showing the lowest modification levels (Fig. 5c-d). All three A3 substrates were efficiently mono-ubiquitinated by the E2 conjugase UBE2A (a.k.a. RAD6A) in the absence of its E3 ligase UBR4 (Fig. 5e). Addition of UBR4 efficiently extended the mono-ubiquitin into poly-ubiquitin chains (Fig. 5e; compare samples 4-6 with 7-9). These data suggest that UBR5 and HUWE1 indeed poly-ubiquitinate A3H independently. In contrast, UBR4 requires prior substrate mono-ubiquitination by its specific E2 conjugase UBE2A and adopts an E4 ligase poly-ubiquitin chain-extending function. However, UBR4-dependent chain extension is not haplotype-specific, suggesting that UBR4 can likely extend any pre-formed chains on A3H.

Cell-based assays had shown that while A3H-I is turned-over by the proteasome, engineered RNA-binding mutants were substantially more unstable (Fig. 4a-b). Since A3H-I purified from *E. coli* bound to substantial amounts of bacterial RNA, we speculated that it might bind RNA in cells. However, the equilibrium may be shifted to a larger unbound fraction of A3H-I in cells compared to A3H-II. Based on the SEC profiles, we predicted that the vast majority of purified WT A3H protein molecules were RNA-bound in our preparations (Fig. S5c), possibly not fully mirroring in-cell equilibrium conditions. Consistent with this notion, RNA-bound A3H-I and A3H-II were substantially less ubiquitinated *in vitro* by each of the three E3 ligases compared to A3H-II-RBM. We hypothesized that this difference might stem from the co-purified RNA shielding the A3H substrates from E3 ligase engagement and predicted that RNA removal during the reaction would render them susceptible to ubiquitination. To test this, RNase A was added to the ubiquitination reaction mixtures. Under these conditions, both A3H-I and A3H-II were ubiquitinated by UBR5, HUWE1, and UBR4 to a similar extent as the A3H-II RBM (Fig. 5f-h). In contrast, ubiquitination of the RNA-free A3H-II-RBM mutant was independent of RNase A addition. From these data, we concluded that RNA indeed shields either the ubiquitination-or E3 ligase recognition site of A3H. Given that ubiquitination of multiple lysine residues contributes to A3H turnover (Fig. 1g-h), we concluded that the RNA-binding region in A3H is likely the recognition site of the E3 ligases, after which the flexibility of these large E3 ligases would allow ubiquitination of multiple surface lysine residues.

Together, these data showed that RNA binding ‘shields’ A3H from ubiquitination by the E3 ligases. In the absence of RNA binding, UBR5 and HUWE1 independently poly-ubiquitinate A3H. Similarly, in the absence of RNA binding, mono-ubiquitination by UBE2A was enhanced, which further increased UBR4 E4 ligase activity, extending poly-ubiquitin chains on pre-formed (mono)-ubiquitin marks (Fig. 5h). Based on these findings and published data^57^, it seems likely that UBR4 can additionally amplify poly-ubiquitination in cellular contexts, by targeting substrates partially ubiquitinated by UBR5 or HUWE1, or factors that were not identified in our screens.

### E3 ligase loss or mutation increases APOBEC signature mutations

Single Base Substitution (SBS) signature mutations are specific patterns of genetic mutations, categorized based on their types and surrounding sequence context. Unbiased computational analysis has identified over 30 different SBS patterns; among these are two highly APOBEC-specific signatures (SBS2, SBS13)^24,30^.

As ablation of *UBR4*, *UBR5*, and *HUWE1* elevated levels of cancer-associated A3B and A3H-I (Fig. 2e-h), we hypothesized that this would increase A3-dependent mutagenesis. To test this hypothesis in an experimentally feasible time frame, we generated an RKO cell line with exogenous A3H-I-driven mutagenic activity (Fig. S6a) and analyzed APOBEC signature mutations. RKO cells have no detectable endogenous A3A or A3H expression, yet express detectable A3B protein levels (Fig. 2g). Therefore, these cells would predominantly accumulate APOBEC mutations driven by the exogenous A3H-I, and potentially to a lesser extent, by endogenous A3B.

These cells were transduced with sgRNAs individually targeting *UBR4*, *UBR5*, or *HUWE1*, sorted for sgRNA-positive cells, after which gene-editing was induced with DOX. Given some eventual growth impairment of E3 ligase knock-out cell pools and potential out-selection over extended cultivation periods, the maximum analysis period with a substantial E3 ligase knock-out population after Cas9 induction was restricted to four doublings (Fig. S6b). Cellular gDNA was isolated from these samples and mutREAD sequencing was performed to determine changes in SBS signature mutations in the gDNA of the analyzed cell pools in an unbiased manner (Fig. 6a)^24,30,58^. This analysis yields a relative distribution of all SBS signature mutations identified in the gDNA of the different genotypes.

**Figure 6.**
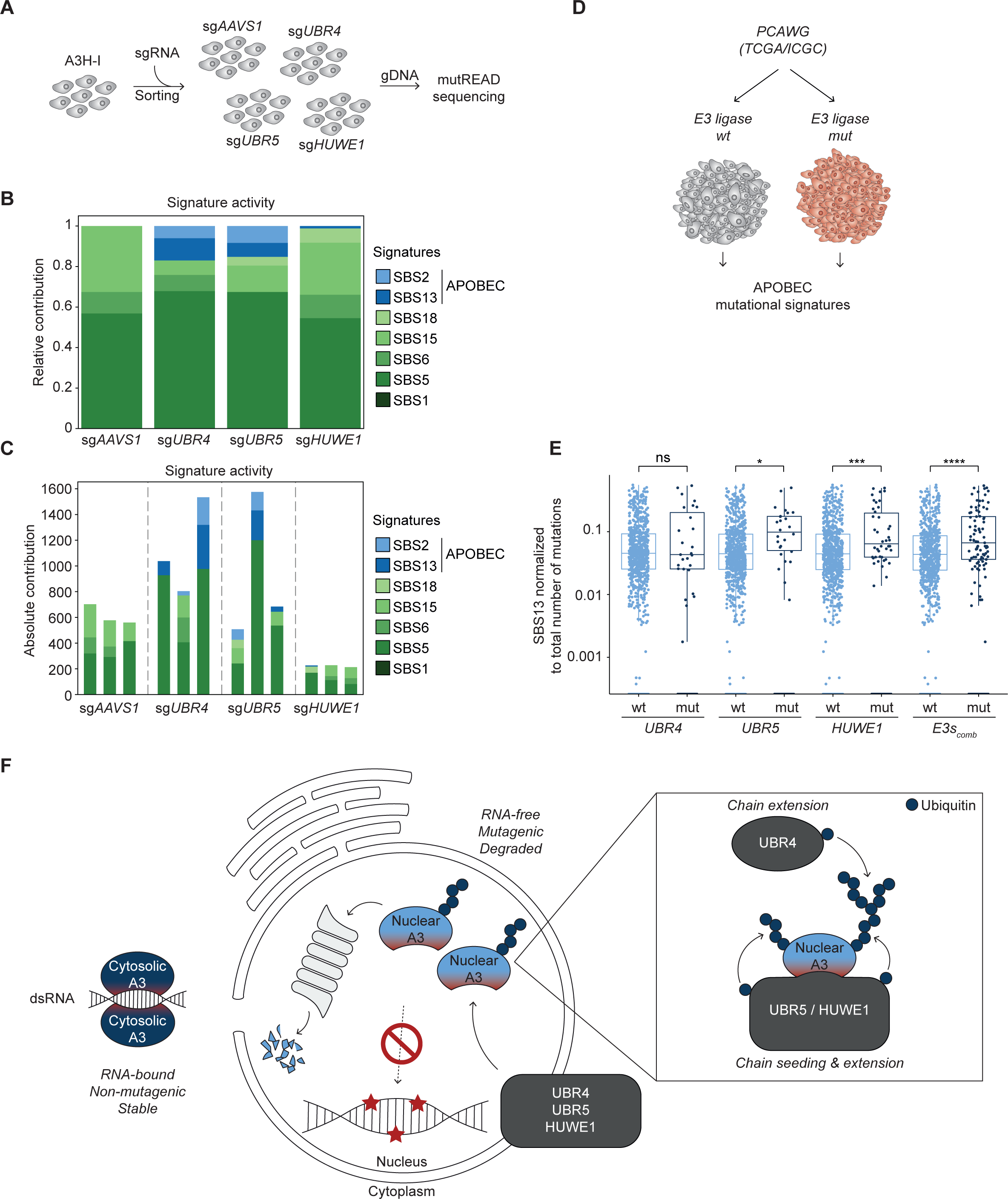
E3 ligase loss or mutation increases APOBEC signature mutations. **(a)** Schematic representation of mutREAD sequencing to identify APOBEC signature mutations in cells. A *UNG2*^-/-^ RKO cell line expressing DOX-inducible Cas9 and mCherry-P2A-3xHA-A3H-I was transduced with sgRNAs targeting *UBR4*, *UBR5*, or *HUWE1*, and subsequently sorted for sgRNA-positive cells. Gene editing was induced with DOX for up to four cell doublings, after which gDNA was isolated and samples prepared for mutREAD sequencing. **(b-c)** Best subset signature refitting of mutREAD data, averaged per genotype, using signatures related to overactivity of APOBEC family enzymes and signatures commonly active in colon carcinoma, the parent tumor type for the model cell line RKO. For the purpose of comparing signature activity across genotypes, samples were matched by recorded doubling time. Each bar represents the average signature refitting results for 3 technical replicates, **(b)** scaled to 1 to show the relative signature contribution, or **(c)** unscaled to show the absolute signature contribution in terms of number of mutations for individual replicates. **(d)** Schematic representation of mutational signature analysis from PCAWG dataset. **(e)** Cancers from TCGA/ICGC were grouped, based on whether the indicated E3 ligase genes were wild-type (wt) or mutated (mut). Mutational signatures were normalized to the total number of mutations in each sample. The level of SBS13 APOBEC signature was compared between the two groups. “E3s_comb_” compares all samples, in which *UBR4*, *UBR5*, and *HUWE1* are either all wild-type, or at least one of the E3 ligase genes was mutated. Data represent Wilcox rank sum test between wt and mut of each genotype, ns: p > 0.05, *: p ≤ 0.05, **: p ≤ 0.01, ***: p ≤ 0.001, ****: p ≤ 0.0001, n = 2703 cancer genome samples). **(f)** Model of A3 protein abundance regulation. RNA binding shields A3 proteins from UBR5/HUWE1 binding and degradation, ensuring high antiviral A3 protein levels in the cytosol. Lack of RNA binding enables UBR5/HUWE1 binding to the unengaged interaction interface (red), resulting in A3 degradation. Since unengaged A3s localize to the nucleus, this ensures low nuclear A3 levels, thereby protecting from genomic DNA mutagenesis. UBR4 acts as an E4 ligase to extend poly-ubiquitin chains.

In *AAVS1*-targeted control cells, three major SBS signatures were identified (Fig. 6b; SBS5/6/15), commonly found in colorectal cancer cell lines. However, APOBEC-specific signatures were not present in these samples. In contrast, APOBEC signature mutations (SBS2 and SBS13) were readily detected in samples from *UBR4-* and *UBR5-*targeted cells (Fig. 6b-c and S6c-d). The total number of mutations accumulated over the cultivation period increased in sg*UBR4* and sg*UBR5* cells, indicating that the increase in APOBEC signature largely represented elevated mutational activity in these samples (Fig. 6c).

In contrast to the other two E3 genotypes, the increase in APOBEC signature was not as pronounced in *HUWE1* knock-out cells (Fig. 6b-c and S6c-d). This correlated with having fewer mutations (Fig. 6c), and the slowest doubling-time of the tested genotypes. We speculate that cells with the strongest increase in mutagenesis in a less-fit *HUWE1* knock-out background, may have been counter-selected during the cultivation period, resulting in lower mutation rates overall.

Taken together, our data showed that under conditions in which degradation of A3 proteins is compromised, A3-driven signature mutations increase, resulting in an elevated mutational burden on the gDNA.

Lastly, we asked whether our cell-based findings would translate to a human setting. To this end, we tested whether mutation – and thus possible loss of function - of the identified E3 ligases, correlated with a specific increase in APOBEC-signature burden in cancer patient samples. For this purpose, cancer whole-genome sequence samples from the PCAWG study^30,59^ were sorted into bins, in which the gDNA sequence for a particular E3 ligase was either WT (wt) or had at least one SNV or InDel outside of intronic regions (mut) (Fig. 6d). In addition, samples in which any of the three E3 ligase genes (*UBR4*, *UBR5*, *HUWE1*) was mutated (E3s_comb_) were compared to all remaining samples.

Subsequently, SBS mutational signatures were retrieved, and the binned groups of each E3 ligase gene compared in a pairwise fashion. To account for a bias in the total mutational burden in the mutant groups of *UBR4*, *UBR5*, *HUWE1,* and unrelated E3 ligases (Fig. S6e), mutational counts were normalized to the total number of mutations in each individual sample. This enabled direct comparison of relative SBS signature distributions between sample groups.

In line with our cell-based experiments, *UBR5-* and *HUWE1*-mutated cancer samples had significantly more APOBEC signature mutations (SBS2/SBS13) compared to their respective WT control groups (Fig. 6e and S6f). In contrast to our cell-based data, there was no significant difference in APOBEC signature mutations between wild-type and *UBR4*-mutant samples (Fig. 6e and Fig. S6f).

The effect in *UBR5*- and *HUWE1*-mutant samples was specific for APOBEC mutational signatures, as mutations in unrelated E3 ligases did not correlate with significant changes in APOBEC signature mutations (Fig. S6g-h). In addition, mutations in *UBR5* or *HUWE1* did not correlate with an increase in other abundant, non-APOBEC signatures (Fig. S6i-k; SBS1/SBS5/SBS18). In fact, the frequencies of these non-APOBEC signatures were predominantly decreased in mutant samples (Fig. S6i-k), which likely stems from the normalization to total mutation counts (Fig. S6e). This indicates that-as one would expect-there is a bias toward greater numbers of total mutations in mutated samples, yet a significantly higher fraction of APOBEC signatures specifically occur in *UBR5* and *HUWE1* mutant cancers (Fig. 6e and Fig. S6f). In agreement with our cell-based data, this suggests that *UBR5* and *HUWE1* are important for curtailing the levels of specific APOBEC proteins in human cancers thereby limiting APOBEC-driven mutagenesis.

Collectively, this study identifies a hitherto unknown framework of cellular guardians that keeps steady-state levels of nuclear A3 proteins low. Our data support a model in which the core of this network consists of UBR5 and HUWE1, E3 ligases that directly recognize A3B and A3H-I and mark them for degradation (Fig. 6f). UBR4 may play an indirect, amplifying role through its E4 ubiquitin chain extension activity.

UBR5 and HUWE1 specifically engage non-RNA-bound A3 proteins through recognition of their unoccupied RNA-binding surfaces. Since RNA binding is important for A3 cytosolic retention, this framework ensures **i)** specific targeting of nuclear A3 proteins that pose a risk to our genome, while **ii)** leaving cytosolic A3 family members available as antiviral restriction factors. Since broad mutational landscapes in cancers enable escape from therapeutic interventions, our findings enable future studies into the identified E3 ligases as possible diagnostic or therapeutic targets.

## Discussion

High A3 levels have been identified in many cancers, and are correlated with higher A3-induced mutational rates (reviewed in ^60^), with A3A, A3B and A3H-I being the major sources of these signatures^18,28,34,35^. The prevalence of APOBEC signatures, found in >50% of all cancers and across cancer types^30^, underpins the critical importance of understanding how human cells restrict APOBEC-induced mutagenesis of their own genomes. Thus far, studies have focused on the differential transcriptional regulation of cancer-associated A3s to explain the high prevalence of APOBEC signatures in tumors. However, other major modes of A3 regulation may play a critical role in cancer mutagenesis.

Here, we identified a novel post-translational regulatory mechanism regulating cellular A3B and A3H-I protein abundance, two major APOBEC signature drivers^18,28,34,35^. The mechanisms we uncovered limit APOBEC-induced mutagenesis in cells. We show that the loss of the central E3 ligases, UBR4, UBR5, and HUWE1, stabilizes A3B and A3H-I proteins, resulting in their accumulation and eventually increasing APOBEC-related mutational burden. Our findings reveal a previously unknown layer of regulation acting to limit cellular A3 protein levels. In the absence of these guardian factors, A3s may broaden the mutational landscape in late-stage cancer, and affect the development of therapy resistance^33,61–64^.

### RNA binding stabilizes A3 proteins

RNA binding mediates important A3 regulatory functions, bridging A3s to form high-molecular-weight complexes in the cytoplasm and inhibiting catalytic activity^36,51,65–70^. In an antiviral context, this leads to inhibition of A3 deaminase activity by the viral RNA until it is released during reverse transcription. However, nuclear A3 accumulation is often the result of diminished cytosolic RNA binding^39^, and comes at the risk of accumulation of an active enzyme with access to the host genome. This requires a cellular framework to ensure high levels of cytosolic antiviral A3 variants, while limiting the concentrations of nuclear, active A3 variants in order to preserve genome integrity.

Our data provide a mechanistic explanation of how such selectivity is achieved: E3 ligases target the A3 RNA-binding surface, a region that also determines its subcellular localization based on whether RNA is bound. This ensures that variants that bind less RNA and are more nuclear, simultaneously become E3 ligase substrates, resulting in their degradation. Our *in vitro* experiments showed that RNA binding plays a major mechanistic role in determining whether UBR5 and HUWE1 can engage A3 proteins as substrates (Fig. 5). Cell-based microscopy and fractionation experiments showed that A3H-I and RNA-binding mutants are predominantly nuclear and degraded in the nucleus (Fig. 4). This degradation may be mediated by the large fraction of cellular proteasomes residing in the nucleus^44^. However, it remains to be determined where in cells E3-substrate interaction and ubiquitination takes place. Localization of the E3 ligases varies in different cell types, and previous analysis places all three E3 ligases in the nucleus and cytosol^71–78^.

Previous work showed that A3H-II has strong RNA-binding activity^36,51,65–70^. However, since the R105G mutation lies outside of the RNA-binding interface, it was unclear whether A3H-I has differential RNA-binding ability, and whether this may underlie the phenotypic changes in localization and stability. Our cell-based data consistently showed intermediate nuclear localization and instability phenotypes for A3H-I, in between A3H-II and designed RNA-binding mutants. Moreover, SEC analysis of recombinant A3H-I revealed that it still co-purified with substantial amounts of bound RNA, albeit less than its A3H-II counterpart. Moreover, while A3H-II purified in part as RNA-bound dimers, A3H-I was exclusively obtained as an RNA-bound monomer.

In line with the findings of a recent interaction study^79^, our cell-based TurboID proteomics showed that A3H-II and A3H-I share many interactors related to RNA processes, albeit fewer for A3H-I. This further supports a model in which A3H-I is still bound by RNA in cells, but to a lesser extent than A3H-II. Together, our findings indicate that A3H-I is likely a partial RNA-binding mutant, for which the equilibrium in cells is shifted towards a more unbound state. As a consequence, it may be less amenable to forming high molecular weight RNA/A3 RNP complexes, as previously described^36,51,65–70^, allowing passive diffusion into the nucleus^42^. The R105G mutation and structural changes in its β-sheet may affect proper positioning of the distal RNA-binding domain. Nevertheless, we cannot rule out additional effects on protein-protein interactions required for high molecular weight complex formation as seen for A3H-II^36,51,65–70^. At least two nuclear and unstable A3H haplotypes arose independently in evolution^10,12,13,40^, suggesting either negative counter-selection of cytosolic variants, or positive selection for nuclear A3 enzymes. In line with this latter notion, various A3 members have been reported to counter infections of unrelated nuclear viruses and transposable elements with DNA genomes or replication intermediates^80–86^. It is, therefore, essential to balance required nuclear antiviral activity with preventing genomic hypermutation. The identified E3 ligase machinery ensures this balance, by only recognizing nuclear A3s that are not engaged with RNA.

### Substrate preference and cooperativity of UBR4, UBR5, and HUWE1

The three identified A3-targeting E3 ligases, UBR5, HUWE1, and UBR4, have previously been linked to the turnover of a diverse array of substrates, thereby exerting control over numerous cellular processes^57,71,72,74,87–105^. This suggests that these E3 ligases possess the ability to identify multiple substrate classes based on broad biochemical or biophysical characteristics. Supporting this idea, recent structural studies of HUWE1 have revealed three distinct substrate-binding domains^106,107^, facilitating the recognition and ubiquitination of unbound nucleic acid binding proteins as well as ubiquitinated/PARylated substrates. Similarly, UBR5 was recently shown to bind and ubiquitinate unengaged transcription factors^87^, and has the ability to function as a ubiquitin chain elongating E4^90,91^. In combination with our findings in this study, these results indicate that HUWE1 and UBR5 are both important players in recognizing and degrading unengaged DNA- and RNA-binding proteins.

Although comprehensive structural information is currently unavailable for UBR4, its involvement in ubiquitinating aggregation-prone nascent polypeptides during proteotoxic stress^94^, diverse mitochondrial proteins^87,95^, and ER-associated degradation substrates^96^ identified in independent screens suggests that this large E3 ligase may also possess the capability to recognize various substrate classes, potentially contingent upon its interaction with different partners. While UBR4 may define substrate selection in other cellular contexts, our data show that in the context of A3 degradation, it likely plays a role as a chain extending E4 ligase. In line with this, UBR4 was neither enriched in our TurboID experiments, nor identified by co-IP (Fig. 3), pointing towards a more distal engagement. However, it could be that UBR4 is specifically recruited to UBR5-or HUWE1-containing complexes in cells to amplify their ubiquitination, and, thereby, proteasomal degradation. In agreement with previous reports^57,108^, our cell-based (Fig. 2g) and *in vitro* (Fig. 5) data suggest that UBR4, UBR5 and HUWE1 have redundant functions and cooperate to assemble ubiquitin chains on their substrates.

### Maintenance of genome integrity

Prior studies have indicated that A3A and A3B are responsible for a substantial part of APOBEC signatures in cancer^28,34^. The presence of the distinct A3H-I haplotype was linked to increased APOBEC signatures in breast and lung cancers^18,32^. However, additional evidence for a contribution of A3H-I to APOBEC3 signatures in other cancers is sparse and little research focuses on A3H-I. We show that APOBEC signatures accumulate in cells in which A3H-I is exogenously expressed, and that impairment of A3H-I and A3B degradation mediated by UBR4, UBR5 and HUWE1 results in an increase in their protein levels, paralleled by an increase in APOBEC signature mutations (Fig. 6). In agreement, we found a significant correlation between *UBR5* or *HUWE1* mutations in human cancer genomes and increased APOBEC signature mutations in these samples. Therefore, analysis of A3 protein levels and/or the mutational status of UBR5 and HUWE1 may prove helpful as a future diagnostic tool that acts as a proxy for the tumor mutational landscape and indicates the likelihood of developing treatment-resistance.

Several proteasome inhibitors are used as therapy for treating multiple myeloma and mantle cell lymphoma, yet relapses and acquired resistance are frequent^109^. The role of APOBEC mutagenesis in cancer therapy and evolution has been suggested to be a double-edged sword. Depending on the level of APOBEC mutagenesis, it could either contribute to greater treatment effectiveness by driving error-catastrophe and synthetic-lethality in cancer cells, while on the other hand it could have detrimental effects by broadening the mutational landscape in tumors, thereby increasing the frequencies of therapeutic resistance^110^. With our finding that proteasomal degradation plays an important role in regulating cancer-associated APOBEC3 protein levels, the application of proteasome inhibitors in cancer therapy and the resulting increase in cellular A3 levels should be assessed accordingly.

### Future directions

Our data presents the importance of a previously underrated RNA-dependent regulatory mechanism of APOBEC3 protein activity and localization through proteasomal degradation controlling levels of nuclear APOBEC3s. While A3A is even more unstable than A3B and A3H-I, it is not ubiquitinated (Fig. 1a and S1h), indicating that cancer-associated A3A protein levels are controlled by a different cellular mechanism. The lack of ubiquitination could suggest A3A is degraded in a ubiquitin-independent manner. Moreover, previous work has shown that A3A transcription is influenced by proteasome inhibition^111^, possibly contributing to the observed increase in A3A protein levels.

Nevertheless, it raises the question of why A3A is not a UBR5/HUWE1 substrate, in contrast to A3B and A3H-I. A3A differs from A3B and A3H-I as it does not form large multimeric complexes^112–114^. The protein sequence of single-domain A3A resembles the C-terminal domain (CTD) of A3B and A3G. However, only the inactive N-terminal domain (NTD) of dual-domain A3s is responsible for RNA binding^115,116^, and contains the critical RNA-binding residues comprising bulky hydrophobic and positively charged amino acids in loop7 and the α6-helix. As the sequence of A3H is similar to the NTD of A3B, we speculate that A3A does not possess the “RNA-binding surface” and can therefore not be recognized by the E3 ligases identified in this study. Thus, future studies will be necessary to identify mechanisms through which A3A protein levels are regulated by proteasomal degradation.

In sum, our current data identify a critical mechanism by which A3B and A3H-I are linked to APOBEC mutation signatures, making these cancer-associated A3s a strong starting point for future development of therapeutic and diagnostic tools.

## Acknowledgements

Next Generation Sequencing analysis was performed by the Vienna Biocenter Core Facilities using the VBCF instrument pool. Proteomics analyses were performed by the Mass Spectrometry Facility at Max Perutz Labs using the VBCF instrument pool; we particularly thank Markus Hartl and WeiQiang Chen for their expert support. Flow cytometry analyses were performed at the BioOptics FACS Facility at the Max Perutz Labs using the Max Perutz Labs instrument pool; we particularly acknowledge Kitti Csalyi, Thomas Sauer, and Johanna Stranner for expert support. Microscopy was performed at the BioOptics Light Microscopy Facility at the Max Perutz Labs; we thank Thomas Peterbauer and Irmgard Fischer for their expert support and training. We thank Johannes Bock for establishing TurboID-related reagents and methodology, Robert Kurzbauer for purification of recombinant proteins, Anna Hakobyan for advice on cancer genome data analysis, Joanna Loizou for expert advice on DNA damage assays, and Marcel Ooms for APOBEC expertise, discussions, and manuscript feedback. We are grateful to the ‘Signaling Mechanisms in Cellular Homeostasis’ doctoral program community, in particular Thomas Decker, Pavel Kovarik and their labs for their technical expertise and help. We thank Life Science Editors for editing services. The results shown here are in whole or part based upon data generated by the TCGA Research Network: https://www.cancer.gov/tcga

Funding sources: This work was funded by Stand-Alone grants (P30231-B, P30415-B, P36572, P36945), Special Research Grant (SFB grant F79), and Doctoral School grant (DK grant W1261) from the Austrian Science Fund (FWF) to GAV, an Austrian Science Fund Special Research Grant (FWF, SFB F79) to TC, and an ERC European Union’s Horizon 2020 research and innovation program grant (AdG 694978) to TC and the Austrian Science Fund Special Research Grant (SFB grant F79) to GEK. VB and S.Sci are the recipients of a DOC fellowship of the Austrian Academy of Sciences. Research at the IMP is supported by Boehringer Ingelheim and the Austrian Research Promotion Agency (Headquarter grant FFG-852936).

## Materials availability statement

All data generated or analyzed during this study are included in the manuscript and supporting files. Data and code pertaining to genomics and mutational signature analysis can be found on https://github.com/menchelab/apobex.

## Author contributions

Conceptualization, G.A.V., I.S., V.B., D.H., T.C. and E.K.; Methodology, G.A.V., I.S., V.B., M.M., H.H., Z.H., D.B.G. and J.F.E.; Software, M.M, I.S. and S.S.; Validation, I.S. and V.B.; Formal Analysis, I.S., V.B. and M.M.; Investigation, I.S., V.B. and K.H.; Resources, Z.H., D.B.G., J.F.E., S.Sci.; Data Curation, I.S., V.B., M.M and G.A.V.; Writing – Original Draft, I.S., V.B. and G.A.V.; Writing – Review & Editing, I.S., V.B., M.M., H.H., K.H., S.S., Z.H., D.B.G., J.F.E., S.Sci, D.H., J.M., T.C., E.K. and G.A.V.; Visualization, I.S., V.B. and M.M.; Supervision G.A.V.; Project Administration, G.A.V.; Funding Acquisition, G.A.V.

## Materials and methods

### Vectors

The lentiviral human ubiquitin-focused sgRNA library consists of 6 sgRNAs per gene for ubiquitin-proteasome system- and autophagy-related genes, and has been described^117^. Lentiviral vectors expressing single or dual sgRNAs from a U6 promoter and eBFP2 or iRFP from a PGK promoter have been described^44^. sgRNA CDSs were cloned in pLentiv2-U6-PGK-iRFP670-P2A-Neo^44^ to perform knock-outs in RKO cell lines. The Dual-A3H-reporter (pLX303-SFFV-MYC-mCherry-A3H-II-P2A-OLLAS-EGFP-A3H-I) was constructed by cloning the ORF of human A3H-I or A3H-II into a modified pLX303 vector^46,118^. cDNAs encoding A3H-I, A3H-II, A3H-I-G105R, A3H-II-R105-G, A3H-II-W115A, A3H-II-R175/176E, A3H-II-E56A-W115A-R175/176E, A3H-I-K-mutants, A3A, A3C, A3D, A3F, A3G, A3G-RNA-binding mutants were synthesized by Twist Bioscience, or generated by fusion PCR and cloned into a modified pLX303 vector for mammalian expression, or into a modified pET47 containing a decahistidine (10×His) tag followed by a Maltose binding protein (MBP) tag and a 3C site vector for bacterial expression. The plasmids and sgRNAs used in this study are listed in the Key resource table.

### Cell culture

Unless otherwise indicated, experiments in this study were reproduced at least twice in independent experiments. All cell lines used in this study were maintained at 37 °C with 5% CO2 in a humidified incubator, routinely tested for mycoplasma contamination, and authenticated by STR analysis. Human RKO (sex unspecified; American Type Culture Collection (ATCC) cat. no. CRL-2577, RRID:CVCL_0504) and THP-1 cells (male; ATCC cat no. TIB-202, RRID:CVCL_0006) were cultured in RPMI 1640 (Gibco) supplemented with 10% FBS (Sigma-Aldrich), L-glutamine (4 mM, Gibco), sodium pyruvate (1 mM, Sigma-Aldrich) and penicillin/streptomycin (100 U/ml/100 μg/ml, Sigma-Aldrich). HeLa cells (female; ATCC cat. no. CCL-2, RRID:CVCL_0030), Lenti-293T lentiviral packaging cells (female, Clontech, cat. No. 632180), and HEK-293T cells (female; ATCC cat. No. CRL-3216, RRID: RRID:CVCL_0063) were cultured in Dulbecco’s modified Eagle’s medium (DMEM; Sigma-Aldrich) supplemented with 10% FBS and penicillin/streptomycin (Sigma-Aldrich). All cell lines used in this study are listed in the Key resource table.

### Generation of clonal iCas9 cell lines

THP-1-DOX-Cas9-P2A-GFP cells were generated by transducing THP-1 cells with the pRRL-TRE3G-Cas9-P2A-GFP-PGK-IRES-rtTA3 lentiviral vector^46^. Cas9 expression was induced with 500 ng/ml of Doxycycline hyclate (DOX, Sigma-Aldrich) and single cells were sorted by FACS into 96-well plates using a FACSAria III cell sorter (BD Biosciences) to obtain single-cell-derived clones. To generate the genetic screening cell line (Dual-A3H-reporter), RKO-DOX-Cas9-P2A-BFP cells^117^ were transduced with pLX303-SFFV-MYC-mCherry-A3H-II-P2A-OLLAS-EGFP-A3H-I and mCherry^+^/GFP^+^ double-positive cells were sorted by FACS into 96-well plates using a FACSAria III cell sorter (BD Biosciences). The mutREAD cell line was generated by co-transducing pLX303-SFFV-MYC-mCherry-P2A-3xHA-A3H-I and DualCRISPR-hU6-sg*UNG2-*mU6-sg*UNG2*-Thy1.1-P2A-Neo into RKO-DOX-Cas9-P2A-GFP^44^. Following Cas9 induction with 200 ng/ml DOX for 6 days, live cells were immunostained for the Thy1.1 surface marker. In brief, cells were washed with PBS, and incubated for 15 min.at 4 °C in Human BD Fc Block (BD Biosciences) to inhibit non-specific antibody binding. Cells were then stained with APC anti-rat CD90/mouse CD90.1 (Thy-1.1) Antibody (BioLegend, 1:260) for 20 min. at 4 °C. Following two washes, mCherry^+^/Thy1.1^+^ single cells were sorted into 96-well plates. *UNG2* homozygous knock-out was confirmed by PCR amplification, TA-cloning and Sanger sequencing of the targeted *UNG2* locus. Polyclonal RKO cell lines were obtained by transducing the parental RKO-DOX-Cas9-P2A-BFP or RKO-DOX-Cas9-P2A-GFP cells with the respective lentiviral expression plasmids listed in the Key resource table.

### Transfections

Transfections for analysis by WB were performed by mixing DNA and polyethylenimine (PEI, Polysciences) in a 1:3 ratio (w/w) in DMEM (Sigma-Aldrich) without supplements. Transfection was performed using 1000 ng of total DNA per well. The day before transfection, 2×10^5^ HEK-293T cells were seeded in 6-well clusters in fully supplemented DMEM media. 36 h. after transfection cells were harvested, washed with ice cold PBS and stored at −80 °C until further processing.

### Lentivirus production and transduction

Lenti-293T lentiviral packaging cells were transfected with DNA mixes containing lentiviral transfer plasmids, pCRV1-Gag-Pol^119^ and pHCMV-VSV-G^120^ using polyethylenimine (PEI, Polysciences) in a 1:3 μg DNA/ µl PEI ratio in non-supplemented DMEM. Virus-containing supernatants were clarified of cellular debris by filtration through a 0.45 μm filter. Virus-like particles were directly used after harvesting or stored at −80 °C. Target cells were transduced in the presence of 5 μg/ml polybrene (Sigma-Aldrich).

### FACS-based CRISPR–Cas9 screens

Lentivirus-like particles were used to transduce RKO-DOX-Cas9-mCherry-A3H-II-P2A-EGFP-A3H-I cells (Dual-A3H-reporter) at a multiplicity of infection (MOI) of less than 0.2 TU/cell, and 500 to 1,000-fold library representation. The percentage of library-positive cells was determined after 3 days of transduction by immunostaining for the Thy1.1 surface marker and subsequent flow cytometric analysis. RKO cells with integrated lentiviral vectors were selected with G418 (1 mg/ml, Sigma-Aldrich) for 5 days. After G418 selection, Cas9 genome editing was induced with DOX (350 ng/ml, Sigma-Aldrich) and after 3 days and 6 days, cells were sorted by FACS. Therefore, cells were harvested, washed with PBS, resuspended in supplemented RPMI-1640 and sorted using the FACSAria III cell sorter operated by BD FACSDiva software (v8.0). RKO cells were gated for non-debris, singlets, BFP-positive (Cas9 expression), EGFP-positive, mCherry-positive, and 1-2% of cells with the lowest or highest EGFP or mCherry signals were sorted into PBS. At least 1×10^6^ cells were collected for each population at each time point in three independent experiments. Sorted samples were re-analyzed for purity, pelleted and stored at −80 °C until further processing. Additionally, 10 million cells from an unsorted reference sample corresponding to 1,000-fold library representation were collected on each sorting day and stored at −80 °C until further processing.

### Next-generation sequencing library preparation

NGS libraries of sorted and unsorted control samples were processed as previously described^44^. In brief, isolated genomic DNA was subjected to two-step PCR. The first PCR amplified the integrated sgRNA cassettes, whereas the second PCR introduced the Illumina adapters. Purified PCR products’ size distribution and concentrations were measured using a fragment analyzer (Advanced Analytical Technologies). Equimolar ratios of the obtained libraries were pooled and sequenced on a HiSeq 2500 platform (Illumina). Primers used for library amplification are listed in the Key resource table.

### Analysis of pooled CRISPR screens

The analysis of the genetic screens was carried out as previously described^44^. Three biological replicates of each sort were included in the analysis. In brief, sgRNAs enriched on day 3 and day 6 (post-Cas9 induction) sorted samples were compared to the matching unsorted control populations harvested on the same days using MAGeCK^121^. Hits were selected based on log_2_ fold-change and p-value and grouped by functional categories. To exclude unspecific hits, we selected genes enriched in EGFP-A3H-I^high^ cell populations on day 3 with a log_2_ fold-change >0.45 and adj. p-value <0.05, which were neither enriched in mCherry^high^ on day 3 (log_2_ fold-change >0.45, p-value <0.05) nor in GFP^low^ on day 3 (log_2_ fold-change >0.45, p-value <0.05). Similarly, genes enriched in EGFP-A3H-I^high^ cell populations on day 6 with a log_2_ fold-change >0.6 and adj. p-value <0.05, which were neither enriched in mCherry^high^ on day 3 (log_2_ fold-change >0.45, p-value <0.05) or day 6 (log_2_ fold-change >0.6, p-value <0.05) nor in GFP^low^ on day 6 (log_2_ fold-change >0.6, p-value <0.05) were selected.

### Protein half-life determination

To estimate A3H-I or A3H-II protein half-lives, cells were treated with 200 μg/ml of cycloheximide (CHX, Sigma-Aldrich). At the indicated time points, total protein extracts were generated using 2x Disruption buffer (1.05 M Urea, 0.667 M β-Mercaptoethanol and 0.7% SDS) or RIPA buffer supplemented with 1% SDS and Benzonase (50 mM Tris HCl (pH 7.4), 150 mM NaCl, 1% NP-40, 0.5% Sodium Deoxycholate, 1 mM EDTA, 1% SDS, 1 mM PMSF (Sigma-Aldrich), Benzonase (25 U/ml, Merck) and 1X cOmplete Protease Inhibitor Cocktail (Roche), analyzed by WB, quantified and normalized as indicated. Single exponential decay curves were plotted using GraphPad Prism (v9), from which protein half-lives were calculated.

### Immunofluorescence confocal microscopy

250,000 RKO cells stably expressing OLLAS-A3H-I or OLLAS-A3H-II (RKO-MYC-mCherry-P2A-OLLAS-A3H-I/II) or RKO cells stably expressing EGFP-A3H-I/II/I-G105R/-II-R105G/II-W115A/II-R175/176E were seeded onto coverslips. After 48 hours, cells were fixed with 4% paraformaldehyde (PFA) for 15 min. In case of antibody staining, cells were permeabilized with 0.25% Triton X-100 in PBS for 5 min., followed by blocking of non-specific sites by incubation with 1% BSA for 30 min. at RT. Coverslips were incubated for 1 h. at RT with primary anti-A3H antibody (Novus Biologicals) 1:100 in 1% BSA, followed by incubation with 1:800 anti-rabbit IgG Alexa Fluor 488 (Abcam) secondary antibody, and incubation for 5 min. with 0.4X Hoechst (Thermo Fisher Scientific) in PBS. The coverslips were mounted using ProLong Gold Antifade Mountant (Invitrogen). Images were collected using a Zeiss LSM 980 confocal microscope at 40X magnification.

### Subcellular fractionation

Subcellular fractionation was performed as previously described^122^. In brief, two million cells were washed in 1 ml PBS and lysed in 500 µl ice-cold REAP buffer (0.1% NP-40 in 1x PBS) supplemented with 1 mM PMSF (Sigma-Aldrich), 1X cOmplete Protease Inhibitor Cocktail (Roche) and Benzonase. 240 µl of the lysates were collected as whole cell fractions, the remaining lysate was centrifuged at 3,000 x g for 60 sec. at 4°C. 240 µl of supernatants were collected as cytosolic fractions, after which pellets were washed with 500 µl of REAP buffer, collected by centrifugation at 3,000 x g for 60 sec. at 4°C, and then resuspended in 240 µl of REAP buffer (nuclear fraction). All fractions were subsequently supplemented with 6x Laemmli sample buffer (62.5 mM Tris-HCl (pH 6.8), 5.8% Glycerol, 2% SDS and 1.7% β-Mercaptoethanol), and boiled for 10 min. Equal volumes of fractions were loaded on a 12 % SDS polyacrylamide gel.

### Western blot analysis

24-48 hours post seeding, cells were treated with various inhibitors (CHX (200 µg/ml), MG132 (10 µM), EPOX (10 µM), CQ (50 µM), BafA (400 nM), NH4Cl (20mM), Leu (50 µM)) for the indicated time points or left untreated. Cells were lysed in Disruption buffer or RIPA lysis buffer supplemented with 1% SDS and Benzonase. Lysates were rotated for 30 min. at 4 °C and then centrifuged at 18,500 x g for 10 min. at 4 °C. Supernatants were transferred to new tubes and protein concentrations were determined using the BCA Protein Assay Kit (Thermo Fisher Scientific). Between 20-50 μg of protein per sample was mixed with Laemmli sample buffer and boiled for 10 min. Proteins were loaded on 12% or 15% SDS polyacrylamide gels, based on the molecular weight of the protein of interest, or alternatively 3-8% NuPage gels (Invitrogen) to probe for high molecular weight proteins. Proteins were separated by SDS page using Tris-Glycine (25 mM Tris, 192 mM glycine, 0.1% SDS) or Tris-Acetate (2.5 mM Tricine, 2.5 mM Tris, 0.05% SDS) SDS running buffer, respectively. Proteins were blotted on PVDF or nitrocellulose membranes at 4 °C for 75 min. at 300 mA in Towbin buffer (25 mM Tris pH 8.3, 192 mM glycine and 20% ethanol). Membranes were blocked in 5% BSA in PBS-T for 1 h. at RT, and subsequently incubated with primary antibodies diluted in 5% BSA overnight at 4 °C (ARP10 Antibody (Novus, 1:1000), Anti-APOBEC3B Antibody (Abcam, 1:1000), Anti-MYC antibody (Sigma-Aldrich, 1:5000), HA-Tag (C29F4) Rabbit mAb (Cell Signaling Technology, 1:1000), HA-Tag (6E2) Mouse mAb (Cell Signaling Technology, 1:1000), OLLAS Epitope Tag Antibody (L2) (Novus, 1:4000), LC3B Antibody ((Cell Signaling Technology, 1:1000), Ubiquitin (P4D1) (Santa Cruz Biotechnology, 1:1000), Anti-UBR4/p600 antibody (Abcam, 1:1000), Rabbit anti-EDD1 Antibody (Bethyl, 1:1000), Rabbit anti-Lasu1/Ureb1 Antibody (HUWE1) (Bethyl, 1:1000), Monoclonal Anti-α-Tubulin antibody produced in mouse (Sigma-Aldrich, 1:1000), Lamin A/C Antibody (E-1) (Santa Cruz Biotechnology, 1:1000), Monoclonal Anti-Vinculin antibody (Sigma-Aldrich, 1:1000), Anti-beta Actin antibody (HRP) (Abcam, 1:20000)). The next day, the membranes were washed three times for 5 min. each with PBS-T and incubated with HRP-coupled secondary antibodies in 5% skimmed milk for 1 h. at RT (Anti-rabbit IgG, HRP-linked Antibody (Cell Signaling Technology, 1:3500), Anti-mouse IgG, HRP-linked Antibody (Cell Signaling Technology, 1:3500), Goat-anti-mouse IgG Light Chain HRP (1:5000), Goat Anti-Rat IgG H&L (HRP) (Abcam, 1:50000)) and imaged with the ChemiDoc Imaging System (Bio-Rad). Relative protein levels were quantified using Image Lab (Bio-Rad).

### Co-Immunoprecipitation assays

HEK-293T cells from one confluent 35-mm dish were lysed in 100 μl of Frackelton lysis buffer (10 mM Tris (pH 7.4), 50 mM NaCl, 30 mM Na_4_P_2_O_7_, 50 mM NaF, 2 mM EDTA, 1% Triton X-100, 1 mM DTT, 1 mM PMSF (Sigma-Aldrich), and 1X cOmplete Protease Inhibitor Cocktail (Roche)). Cells were incubated on a rotating wheel at 4 °C for 30 min. and subsequently centrifuged at 20,000 x g at 4 °C for 30 min. The supernatant was transferred to a new tube and 10 μl (10% of the lysate used for the IPs) was collected as input. 500 μg of lysates were incubated overnight at 4 °C on a rotating wheel with anti-HA antibody (Cell Signaling Technology, 1:100). The next day, magnetic beads (Protein A/G Magnetic Beads, Thermo Fisher Scientific) used for anti-HA antibody IPs, were blocked by rotation in 3% BSA in Frackelton Buffer for 1 h. at 4 °C. 25 μl of beads were added to 500 μg of lysates and rotated for 2 h. at 4 °C. Then, the beads were washed five times with 1 ml of Frackelton lysis buffer. Proteins were eluted by boiling in 2X Disruption buffer for 10 min. at 95 °C.

### Immunoprecipitations for ubiquitination

HEK-293T cells from confluent 35-mm dishes were lysed in 100 µl of RIPA buffer with 1% SDS (50 mM Tris-HCl (pH 7.4), 150 mM NaCl, 1% SDS, 0.5% sodium deoxycholate, 1% Triton X-100), supplemented with 40 mM N-Ethylmaleimide, 40 mM iodoacetamide, 25 U/ml Benzonase, 1 mM PMSF (Sigma-Aldrich), and 1X cOmplete Protease Inhibitor Cocktail (Roche). Cells were incubated on a rotating wheel at 4 °C for 30 min. and centrifuged at 20,000 x g at 4 °C for 15 min. Supernatants were transferred to new tubes and 30 µg of the lysates were collected as input. 500 μg of lysates were incubated overnight at 4 °C on a rotating wheel with an anti-HA antibody (Cell Signaling Technology, 1:100). The next day, magnetic beads (Protein A/G Magnetic Beads, Thermo Fisher Scientific) were blocked by rotation in 3% BSA in RIPA Buffer for 1 h. at 4 °C. 25 μl of beads were added to 500 μg of lysates and rotated for 2 h. at 4 °C. Subsequently, beads were washed five times with 1 ml of RIPA buffer, supplemented with 300 mM NaCl. Proteins were eluted by boiling in 2X Disruption buffer for 10 min. at 95 °C.

### RNA isolation, cDNA synthesis, and qPCR

For mRNA half-life determination, cells were treated with actinomycin D (ActD) at a final concentration of 5 µg/ml for the indicated times, at which point total RNA was isolated. The decay of *APOBEC3H* mRNA was assessed by RT-qPCR. RNA from 1×10^6^ cells was isolated using Trizol reagent (Thermo Fisher Scientific) and treated with Turbo DNase (Thermo Fisher Scientific). cDNA was prepared using random hexamer primers and RevertAid Reverse Transcriptase (Thermo Fisher Scientific). Real-time PCR experiments were run on a Mastercycler (Biorad), using the Luna Universal qPCR Master Mix (NEB). Primers for qPCR are listed in the Key resource table.

### TurboID

A3H-I and A3H-II, as well as an EGFP control were cloned into lentiviral plasmid pCW-MYC-TurboID-MCS-PGK-mCherry-P2A-rtTA^46^, for the expression of fusion proteins N-terminally tagged with TurboID. For the TurboID experiment, polyclonal cell lines stably expressing DOX-inducible TurboID-A3H-I, TurboID-A3H-II and TurboID-EGFP were stimulated with DOX for 48 h. to induce the expression of the TurboID fusion constructs. Subsequently, cells were stimulated for 5 h. with 10 µM EPOX to inhibit proteasomal degradation, and finally biotinylation was induced for 15 min. by addition of 500 µM Biotin (Sigma) to the cell culture medium. Subsequently, cells were washed 4 times with ice-cold PBS, prior to lysis in RIPA lysis buffer (50 mM Tris HCl (pH 7.4), 150 mM NaCl, 1% NP-40, 0.5% Sodium Deoxycholate, 1 mM EDTA, 0.1% SDS, 1 mM PMSF (Sigma-Aldrich) and 1X cOmplete Protease Inhibitor Cocktail (Roche). Lysates were rotated for 30 min. at 4 °C, centrifuged at 18,500 x g for 10 min. at 4 °C, and protein concentrations were determined by BCA assay. 1200 µg of protein was incubated overnight rotating at 4°C with 200 ul of Streptavidin beads (Thermo Scientific) which were acetylated with Sulfo-NHS-Acetate beforehand, as described^123^. Beads were washed twice with 1 ml of RIPA buffer, once with 1 ml of 2 M Urea in 10 mM Tris (pH 8), twice with 1 ml RIPA buffer and five times with 50 mM HEPES (pH 7.8). Three technical replicates were subjected to nLC-MS/MS analysis.

### Sample preparation for mass spectrometry analysis

Beads were resuspended in 50 µl 1 M Urea and 50 mM ammonium bicarbonate. Disulfide bonds were reduced with 2 µl of 250 mM dithiothreitol (DTT) for 30 min. at room temperature before adding 2 µl of 500 mM iodoacetamide and incubating for 30 min. at room temperature in the dark. Remaining iodoacetamide was quenched with 1 µl of 250 mM DTT for 10 min. Proteins were digested with 150 ng LysC (mass spectrometry grade, FUJIFILM Wako chemicals) at 25 °C overnight. The supernatant was transferred to a new tube and digested with 150 ng trypsin (Trypsin Gold, Promega) in 1.5 µl 50 mM ammonium bicarbonate at 37 °C for 5 h. The digest was stopped by the addition of trifluoroacetic acid (TFA) to a final concentration of 0.5 %, and the peptides were desalted using C18 Stagetips^124^.

### Liquid chromatography-Mass spectrometry data acquisition

Peptides were separated on an Ultimate 3000 RSLC nano-flow chromatography system (Thermo-Fisher), using a pre-column for sample loading (Acclaim PepMap C18, 2 cm × 0.1 mm, 5 μm, Thermo-Fisher), and a C18 analytical column (Acclaim PepMap C18, 50 cm × 0.75 mm, 2 μm, Thermo-Fisher), applying a segmented linear gradient from 2% to 35% and finally 80% solvent B (80 % acetonitrile, 0.1 % formic acid; solvent A 0.1 % formic acid) at a flow rate of 230 nl/min. over 120 min. Eluting peptides were analyzed on an Exploris 480 Orbitrap mass spectrometer (Thermo Fisher) coupled to the column with a FAIMS pro ion-source (Thermo-Fisher) using coated emitter tips (PepSep, MSWil) with the following settings: The mass spectrometer was operated in DDA mode with two FAIMS compensation voltages (CV) set to −45 or −60 and 1.5 s. cycle time per CV. The survey scans were obtained in a mass range of 350-1500 m/z, at a resolution of 60k at 200 m/z, and a normalized AGC target at 100%. The most intense ions were selected with an isolation width of 1 m/z, fragmented in the HCD cell at 28% collision energy, and the spectra recorded for max. 50 ms. at a normalized AGC target of 100% and a resolution of 15k. Peptides with a charge of +2 to +6 were included for fragmentation, the peptide match feature was set to preferred, the exclude isotope feature was enabled, and selected precursors were dynamically excluded from repeated sampling for 45 s.

### Data analysis

MS raw data split for each CV using FreeStyle 1.7 (Thermo Fisher), were analyzed using the MaxQuant software package (version 1.6.17.0)^125^ with the Uniprot human reference proteome (version 2020_01)^126^, as well as a database of most common contaminants. The search was performed with full trypsin specificity and a maximum of two missed cleavages at a protein and peptide spectrum match false discovery rate of 1%. Carbamidomethylation of cysteine residues was set as fixed, oxidation of methionine, phosphorylation (STY) and N-terminal acetylation as variable modifications. For label-free quantification the “match between runs” only within the sample batch and the LFQ function were activated - all other parameters were left at default. MaxQuant output tables were further processed in R 4.2.1^127^ using Cassiopeia_LFQ^128^. Reverse database identifications, contaminant proteins, protein groups identified only by a modified peptide, protein groups with less than two quantitative values in one experimental group, and protein groups with less than 2 razor peptides were removed for further analysis. Missing values were replaced by randomly drawing data points from a normal distribution model on the whole dataset (data mean shifted by −1.8 standard deviations, a width of the distribution of 0.3 standard deviations). Differential expression analysis was performed with Enrichr^129–131^. In detail, differentially enriched proteins in A3H-I/GFP (light blue, LFC >1, p-value <0.01) and A3H-II/GFP (dark blue, LFC >1, p-value <0.01) were compared using RStudio (v4.1.3). GO:CC and GO: BP terms were derived for proteins being specifically enriched in the A3H-I or A3H-II interactome, or for shared interactors.

### Proteomics data deposition

The mass spectrometry proteomics data have been deposited to the ProteomeXchange Consortium via the PRIDE partner repository^132^ with the dataset identifier PXD051267.

### Protein purification

BL21 (DE3) RIPL *E. coli* cells harboring 10xHis-MBP-A3H expression vectors were grown at 37 °C. IPTG at 0.3 mM was added when OD600 reached about 0.6–0.8 for overnight induction at 18°C. To purify dimer and monomer of 10xHis-MBP-fused A3H, *E. coli* cells expressing MBP-A3H-I/II/II-E56A-W115A-R175/176E were harvested and lysed using an EmulsiFlex C3 homogenizer in buffer A (25 mM HEPES pH 7.5, 500 mM NaCl, 20mM imidazole, 5% glycerol, 5mM β-Mercaptoethanol, 1X EDTA-free cOmplete Protease Inhibitor Cocktail (Roche), 0.5 mM PMSF (Sigma-Aldrich), 100 ug/ml RNase A (Carl Roth)). The clear soluble fraction obtained after centrifugation (45 min. at 18.000 x g) was loaded onto a HisTrap HP 5 mL, washed with wash buffer (25 mM HEPES pH 7.5, 500 mM NaCl, 20mM imidazole, 5% glycerol, 5mM β-Mercaptoethanol) and eluted with buffer B (25 mM HEPES pH 7.5, 500 mM NaCl, 500mM imidazole, 5% glycerol, 5mM β-Mercaptoethanol). Fractions containing A3H were identified by SDS page, concentrated, and separated by Superdex 200 10/300 gel filtration chromatography (Cytiva) in SEC buffer (25 mM HEPES pH 7.5, 250 mM NaCl, 5% glycerol, 2 mM DTT. The dimer and monomer fractions were collected, concentrated and flash frozen in liquid nitrogen. Codon-optimized human HUWE1 was synthesized in fragments and assembled into a GoldenBac pGBdest vector using a BsaI-GoldenGate reaction^133^. The construct had a his_8_ tag followed by a rigid enhancer linker (AEAAAKEAAAKEAAAKEAAAKALEAEAAAKEAAAKEAAAKEAAAKA) and then a TEV cleavage site. Human UBR4 was cloned by the same procedure, except the tag used was a C-terminal Strep tag separated from the protein by a 3C cleavage site. Plasmids were transformed into DH10 MultiBac cells for bacmid generation. *Spodoptera frugiperda* (Sf9) cells in ESF921 serum-free growth medium (Expression Systems) were transfected with the bacmids for virus amplification, which was monitored using yellow fluorescent protein signal. HUWE1 and UBR4 were then expressed in *Trichoplusia ni* High-Five insect cells (Thermo Fisher) with a density of 1.5 x10^6^ using a 1:70 inoculation from the V1 stock for 92 hours at 21°C in Insect Xpress Protein-free Insect Cell Medium (Lonza) supplemented with GlutaMAX (GIBCO) and Pen/Strep Amphotericin B (Lonza). Cells were harvested by centrifugation at 700 x g, washed in PBS and then flash frozen and stored at −70°C. For HUWE1, the thawed pellet was resuspended in 50 mM HEPES pH 7.5, 300 mM NaCl, 0.5 mM TCEP, 20 mM imidazole with Complete EDTA-free Protease inhibitor (Roche) and 20 µl Benzonase (IMP Molecular Biology Service) and lysed using dounce homogenization. The lysate was cleared by centrifugation at 40,000 x g. The soluble fraction was then loaded on a 5 ml HisTrap HP column (Cytiva) pre-equilibrated in the lysis buffer using an Äkta Pure 25 system (Cytiva). The column was washed with 10 column volumes of the same buffer, followed by 7 column volumes of the same buffer but with 35 mM imidazole, and then eluted with 300 mM imidazole. The protein was cleaved with TEV protease and simultaneously dialyzed to remove imidazole overnight. The cleaved protein was then reapplied to the HisTrap HP column in 20 mM imidazole and the flow through was collected and concentrated to 1.5 ml. Finally, the protein was applied to a Superose 6 16/60 column (Cytiva) equilibrated in 50 mM HEPES pH 7.5, 150 mM NaCl, 0.5 mM TCEP. Protein-containing fractions were pooled and concentrated. The protein was pure as assessed by SDS-PAGE and the concentration was estimated by A280 absorption using an extinction coefficient of 251,770 M^-1^ cm^-1^. For UBR4, the same lysis procedure was used except with phosphate buffered saline (PBS), 0.5 mM TCEP at pH 7.4 as the buffer. The soluble lysate was then loaded on a 5 ml StrepTrap HP column (Cytiva) equilibrated in PBS, washed with the same buffer and then eluted with 2.5 mM desthiobiotin. Eluted UBR4 was further applied to a Resource Q column (Cytiva) equilibrated in PBS for further purification by anion exchange chromatography using a 250 to 500 mM NaCl gradient. UBR4-containing fractions were pooled and concentrated and the final concentration measured by absorption at 280 nm using a calculated extinction coefficient of 472,140 M^-1^cm^-1^ The codon-optimized UBE2A gene was synthesized by Twist Bioscience. The NEB HiFi assembly kit (New England Biolabs) was used to clone the gene into a pET29b expression vector, which contained an N-terminal His6-tag and TEV cleavage site. Expression was performed using *E. coli* BL21 (DE3) cells with IPTG induction at 20 °C overnight. Cells were resuspended in buffer containing 50 mM Tris pH 7.5, 150 mM NaCl, 10 mM imidazole, Complete EDTA-free protease inhibitor cocktail. Cells were lysed by sonication and clarified by centrifugation at 18,500*g* for 20 min at 4 °C. Clarified lysates were then incubated with Ni-NTA resin (Qiagen) for 1 h at 4 °C with mild agitation. Ni-NTA resin was washed before elution with 150 mM imidazole. UBE2A was further purified by SEC using a Superdex 75 16/600 column (Cytiva) equilibrated in 50 mM Tris, 150 mM NaCl and 0.5 mM TCEP, pH 7.5. Recombinant human UBR5, Ubiquitin, DyLight488-labeled Ubiquitin, UBE2D3 and UBA1 were purified as described previously^91,107,134^.

### UREA denaturing gel

To determine the amount of bound RNA, equal protein amounts of 10xHis-MBP-A3H-I/II/II-E56A-W115A-R175/176E were denatured by the addition of 2x denaturing buffer (0.6 g/ml urea, 0.1% SDS, 1 mM EDTA pH 8.0, 0.5 mg/ml Xylene Cyanol, 0.5 mg/ml Bromophenol blue) and incubation for 5 min. at 95 °C. Subsequently, the RNA was separated from the proteins on a 6%Urea-TBE gel and visualized by SYBR Gold staining.

### *In vitro* ubiquitination assays

*In vitro* ubiquitination assays contained 10 μM ubiquitin, 1 μM DyLight488-labeled ubiquitin for in-gel visualization, 0.2 μM E1 (UBA1), 0.4 μM E2 (UBE2A for UBR4, UBE2D3 for UBR5 and HUWE1), 0.4 μM E3 (UBR4, UBR5, HUWE1) and 4 μM substrate protein (10xHis-MBP-A3H), unless indicated otherwise. The assays were performed in 25 mM HEPES pH 7.5, 200 mM NaCl, 5 mM MgCl2, 0.5 mM TCEP (assay buffer) at 30 °C (UBR5) or 37°C (UBR4 and HUWE1) for 1 h. in 11 μl volumes. Reactions were initiated by the addition of 5 mM ATP. After one hour, reactions were diluted in 500 µl immune-precipitation buffer (RIPA buffer supplemented with 40 mM N-Ethylmaleimide (Sigma-Aldrich), 40 mM iodoacetamide (Sigma-Aldrich), 1 mM PMSF (Sigma-Aldrich), and 1X cOmplete Protease Inhibitor Cocktail (Roche) and incubated with 15 µl of anti-MBP couped beads (pre-blocked for 1h in RIPA with 3 % BSA; NEB) for 2 h. at 4°C. Beads were washed 5 times with RIPA-300 mM NaCl, followed by elution of A3H in 2x Laemmli sample buffer with 2 mM DTT. SDS-PAGE was performed using 4–20% Mini-PROTEAN TGX Stain-Free (BioRad), and ubiquitinated A3H was detected by visualization of DyLight488-ubiquitin using ChemiDoc MP system (Bio-Rad).

### mutREAD sequencing and detection of mutational signatures

A monoclonal RKO-DOX-Cas9-P2A-GFP cell line expressing MYC-mCherry-P2A-3xHA-A3H-I that also has a homozygous knock-out of *UNG2*^-/-^ was transduced with DualCRISPR-hU6-sgRNA-mU6-sgRNA-iRFP670 targeting indicated genes and bulk sorted for iRFP^+^ cells. Cas9 was induced with DOX and sequential samples collected up to 10 days after induction. Their doubling times were calculated based on cell numbers counted during cell passaging, and samples of similar doublings compared for analysis. Cells were harvested for WB to verify E3 ligase knock-out and gDNA extraction. DNA was isolated using the DNeasy Blood & Tissue Kit (Qiagen) after which samples were subjected to mutREAD library preparation as described^58^. In brief, 500 ng gDNA was digested with 50 U PstI-HF (NEB) and 50 U ApoI-HF (NEB) together with 0.187 µM annealed mutREAD adapters (see Key resource table), 400 U T4 DNA ligase (NEB), 1 mM ATP in 1X CutSmart buffer. The 50 µl reaction was incubated for 3 h. at 30 °C in a thermal cycler and subsequently stopped by addition of 10 µL 50 mM EDTA. DNA fragments were size selected for 400-600 bp fragments performing a 0.6X/0.15X 2-sided size selection using DNA clean beads (VAHTS). Size-selected DNA fragments were amplified using NEB unique multiplexed i5 and i7 primers (E6440) in a total reaction volume of 100 µl together with 2 U Phusion High-Fidelity DNA Polymerase (NEB) and in the presence of 0.2 mM dNTPs and 1X Phusion High-Fidelity buffer. PCR was performed in the following conditions: 98 °C/2 min. denaturation, 12 cycles of amplification at 98 °C/10 s, 65 °C/30 s, 72 °C/30 s and final extension at 72 °C for 5 min. Subsequently, libraries were selected by again performing a 0.6X/0.15X 2-sided size selection after which samples were eluted in 20 μl TE buffer (Tris-EDTA buffer 10 mM Tris-HCl and 0.1 mM EDTA, pH 8) and stored at −20 °C. Quality and fragment sizes were evaluated on a fragment analyzer (Advanced Analytical Technologies) and quantified using Picogreen assay (Invitrogen). Equimolar ratios of the obtained libraries were pooled and 150 bp paired-end sequenced on an Illumina NovaSeq S4 platform.

### Alignment and Somatic Variant Detection

Raw reads were processed according to GATK4 best practices^135^. Briefly, reads were mapped to the GRCh38 human reference genome and single nucleotide substitutions and InDels were called with Mutect2 in tumor-only mode^136^. Resulting variants were filtered by only retaining variants with Mutect2 quality status ‘PASS’ and variant allele frequency (VAF) between 0.3 and 0.7. Variants were further filtered by retaining only variants which originated from reads aligned to chromosomes 1-22, X or Y. In order to exclude cell-line specific variants found in the parental clone, we obtained 30x whole genome sequencing (BGITech Global, BGI Genomics Co., Ltd.) from the parental clone (RKO-DOX-Cas9-P2A-GFP) and retained only variants which were not also present in the parental clone.

### Bootstrapped Background Correction

We applied a bootstrapping strategy to construct an average background mutational profile from the samples obtained 3 days after transduction with the control sgRNA vector targeting *AAVS1* (n = 3). First, to reduce the effect of outliers, the mutational count of all samples was down-scaled proportional to the sample with the lowest mutation count. For each sample, 10,000 bootstrap samples were generated using weighted random multinomial sampling, with the underlying distribution of mutation counts across the 96 features serving as weights. This process yielded 30,000 bootstrapped samples in total, from which the mean mutation count across the 96 mutation channels was computed, resulting in the final average background profile. The average background mutational profile was subsequently subtracted from the mutational matrix of the samples using element-wise subtraction with thresholding at 0.

### Mutational Signature Analysis

We conducted signature refitting and plotting using the MutationalPatterns package^137^ in R. Refitting was accomplished by incorporating a catalog of signatures frequently active in colon carcinoma, alongside APOBEC-related signatures (SBS2 and SBS13), using the best subset refit approach with the parameter max_delta set to 0.001 to obtain a strict estimate of signature activity. The significance of the refitting results was assessed using the signature activity test from the mSigAct package^138^. For the purpose of comparing signature activity across genotypes, samples were matched by the number of cell doublings.

### Computational analysis of cancer genome sequencing database

SBS mutational signatures analyzed in the PCAWG study were obtained from the ICGC Data portal^139^. Corresponding vcf files for each sample were either retrieved from the ICGC Data portal or through granted access of the TCGA Research Network^140^. To identify the genotype of a specific gene of interested (E3 ligase) in each sample, the corresponding vcf files (snv and indel) were tested for variants within the location of the coding gene. Samples exhibiting at least one variant in either snv or indel with a Variant Classification not being “Intron”, were categorized as “mut”. Conversely, the remaining samples were categorized as “wt”. SBS mutational signatures were normalized to the total number of mutations in each sample and the fraction of SBS signatures of interest compared between “wt”’ and “mut” groups. Four E3 ligase genes (*HECTD4*, *NEDD4L*, *HECT1*, *PCHF3*), with a similar number of “mut” cancers were randomly selected and used for probing specificity.

### Quantification and statistical analysis

Sample number and number of biological replicates are indicated in the figure legends of all figures. Statistical analyses were performed with the R (v4.0.2) programming environment using RStudio (v4.1.3) or GraphPad Prism 10.0.2.

---

**Figure S1.**
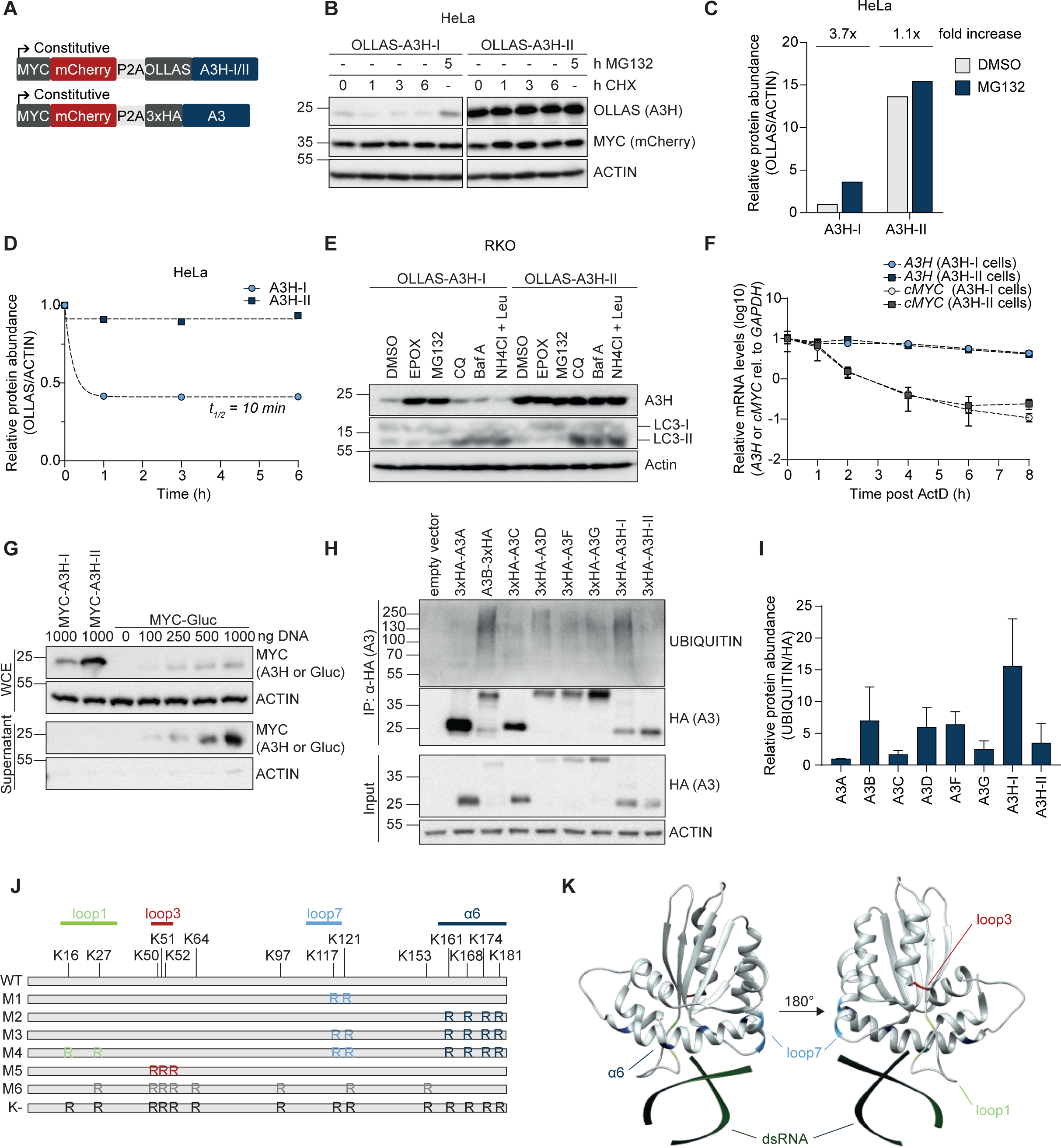
Proteasomal degradation controls protein levels of cancer-associated A3s. **(a)** Schematic representation of the lentiviral mCherry-P2A-OLLAS-A3 and mCherry-P2A-3xHA-A3 expression constructs used in this study. The ribosomal skip site P2A ensures equimolar translation of the stable internal control mCherry and tagged A3 protein. **(b-d)** Lentiviral constructs encoding mCherry-P2A-OLLAS-A3H-I/II were stably integrated in HeLa cells by lentiviral transduction. Polyclonal cell pools were treated with MG132 or CHX for the indicated times, followed by **(b)** analysis of protein levels by WB, **(c)** quantification of relative A3H-I and A3H-II protein levels upon proteasome inhibition, or **(d)** translation inhibition, by calculating single-step exponential decay curves to derive protein half-live. **(e)** RKO cells expressing the indicated OLLAS-tagged-A3H haplotypes were treated with different proteasome inhibitors (EPOX, MG132), or autophagy/lysosomal degradation inhibitors (chloroquine (CQ)), bafilomycin A (BafA), or leupeptin (Leu)) for 5 h. LC3-I to LC3-II conversion was detected as a marker for inhibition of lysosomal degradation. **(f)** RKO cells stably expressing OLLAS-A3H were treated for the indicated times with Actinomycin D (ActD), after which relative exogenous *APOBEC3H* and endogenous *cMYC* mRNA levels were quantified by RT-qPCR. **(g)** HEK-293T cells were transfected with plasmids encoding MYC-tagged A3H-I/II or Gaussia luciferase (MYC-Gluc). 48 h. post transfection, MYC-tagged protein levels in the supernatant and the whole cell extract (WCE) were analyzed by WB. **(h-i)** HEK-293T cells were transfected with different amounts of the indicated 3xHA-A3 plasmids to achieve similar steady-state A3 protein levels. After treatment with EPOX for 5 h., 3xHA-tagged proteins were immunoprecipitated from cell lysates, and their ubiquitination analyzed by **(h)** WB with a total ubiquitin antibody, and **(i)** quantified (means and SD, n = 4). **(j)** Schematic representation of lysine positions in A3H-I. Lysine residues were systematically grouped based on their proximity in the A3H structure and mutated to arginine (M1-M6). In K-, all 14 lysine residues were mutated to arginine. **(k)** The positions of the mutated lysine residues in the A3H-II crystal structure are highlighted in different colors (PDB: 6B0B).

**Figure S2.**
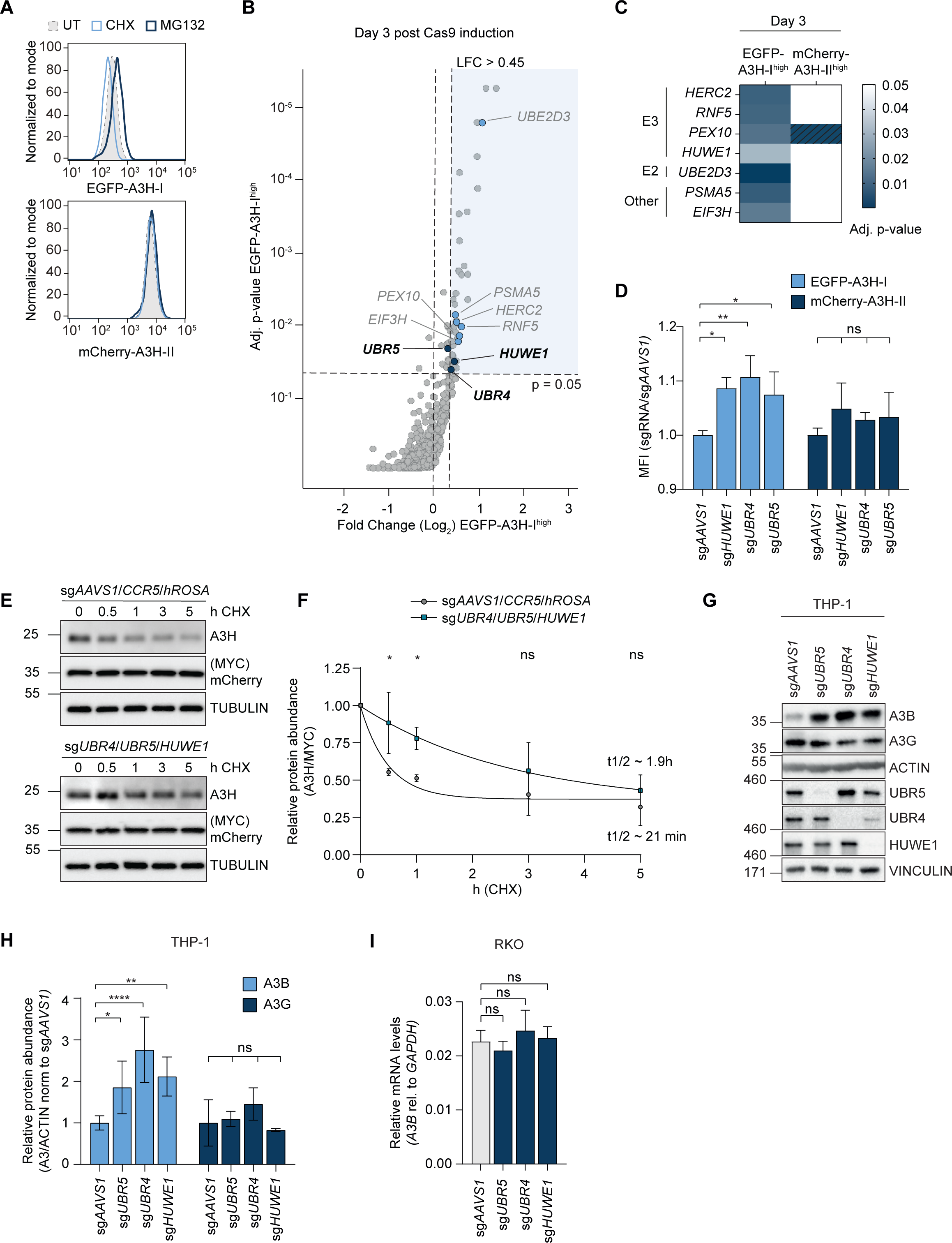
The E3 ligases UBR4, UBR5, and HUWE1 independently mediate turnover of A3B and A3H-I. **(a)** Monoclonal RKO-DOX-Cas9-dualA3H cells were treated with CHX or EPOX for 5 h., followed by analysis of EGFP-A3H-I or mCherry-A3H-II levels by flow cytometry. **(b)** Targeted genes enriched in EGFP-A3H-I^high^ sorted cell populations 3 days post Cas9 induction. **(c)** Heatmap of top genes on log_2_ fold-change and p-value grouped by functional categories. Genes enriched in EGFP-A3H-I^high^ cell populations 3 days post Cas9 induction with a log_2_ fold-change > 0.45 which were not enriched in mCherry^high^ or GFP^low^ on the same day. Dashed lines indicate a log_2_ fold-change < 0.45. Adjusted p-values are based on MaGECK analysis of three independent replicate sorts. E3 (E3 ligases), E2 (E2 conjungating enzymes). **(d)** Polyclonal RKO-DOX-Cas9-dualA3H cells were transduced with sgRNA vectors targeting *UBR4*, *UBR5*, or *HUWE1*. EGFP-A3H-I and mCherry-A3H-II abundance was analyzed by flow cytometry 6 days after Cas9 induction. The mean fluorescence intensity (MFI) was quantified (one-way ANOVA compared to sg*AAVS1*, ** p < 0.005, * p < 0.05, n = 3). **(e)** RKO-DOX-Cas9-MYC-mCherry-P2A-3xHA-A3H-I cells were transduced with sgRNA simultaneously targeting either the three E3 ligases or control loci (*AAVS1*, *CCR5*, *hROSA*), gene editing induced with DOX for 6 days and then treated with CHX for different times. A3H-I protein levels were determined by WB and **(f)** the half-life quantified from single-step exponential decay curves, statistics were calculated by two-way ANOVA, * p < 0.05 (n = 3). **(g)** THP-1 cells harboring DOX-inducible Cas9 were transduced with sgRNAs targeting *UBR4*, *UBR5*, or *HUWE1*, and sorted for sgRNA-positive cells. Gene editing was induced with DOX for 3 days, after which endogenous A3B and A3G protein levels were determined by WB, and **(h)** quantified (means and SD, two-way ANOVA, **** p < 0.0001, *** p < 0.0005, ** p < 0.005, * p < 0.05, n = 4). **(i)** RKO cells harboring DOX-inducible Cas9 were transduced with sgRNAs targeting *UBR4*, *UBR5*, or *HUWE1*, and sorted for sgRNA-positive cells. Gene editing was induced with DOX for 6 days, after which endogenous *A3B* mRNA levels were determined by RT-qPCR (means and SD, one-way ANOVA, ns p > 0.1, n = 3).

**Figure S3.**
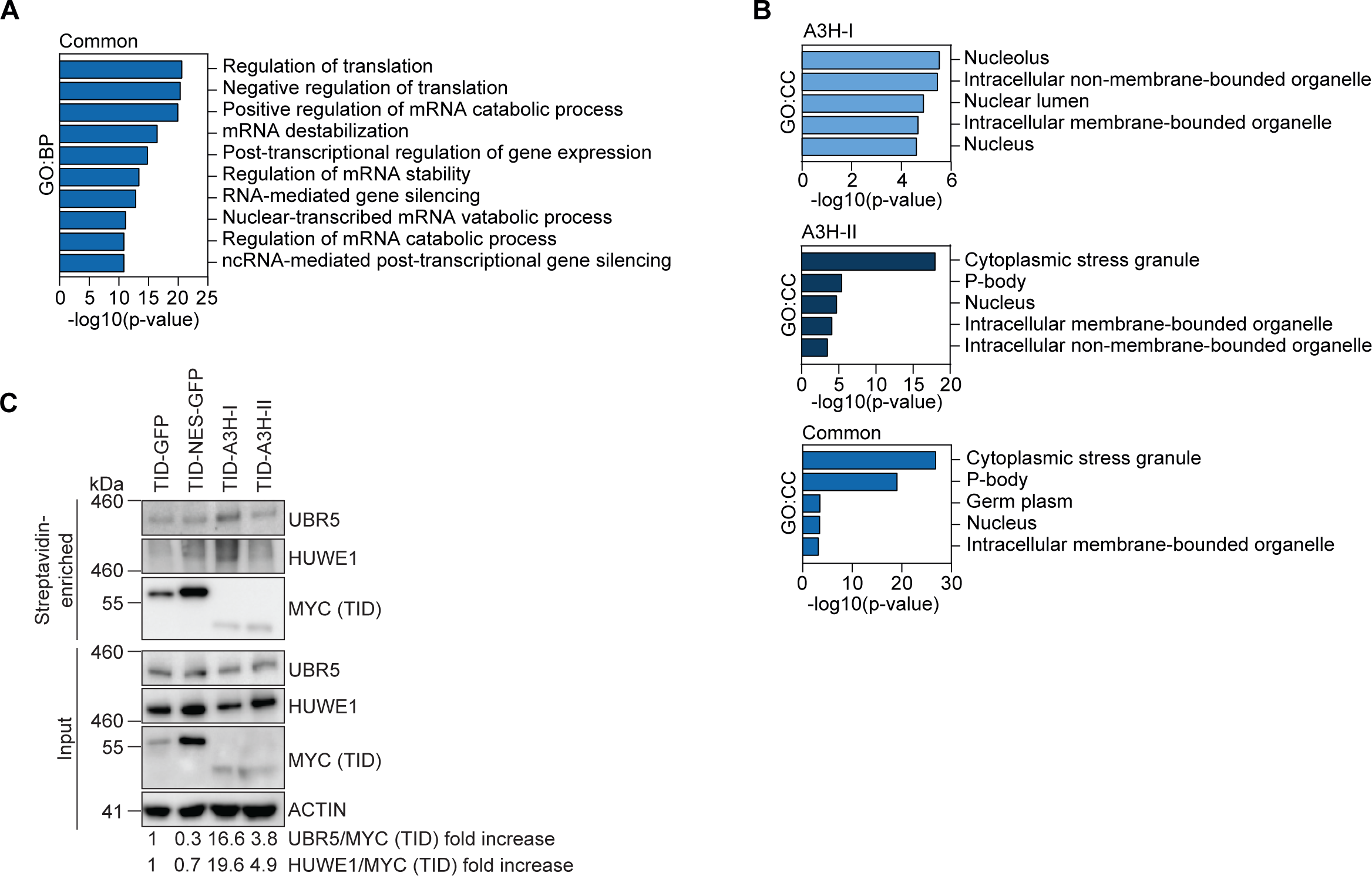
UBR5 and HUWE1 form a complex with A3H-I and other unstable A3 deaminases in cells. **(a)** GO terms for biological processes (GO:BP) of differentially enriched proteins shared between A3H-I/GFP and A3H-II/GFP (LFC > 1, p-value < 0.01, input: 121 factors derived from Fig. 3b) **(b)** GO terms for cellular compartments (GO:CC) of differentially enriched proteins in A3H-I/GFP (light blue, input: 170 factors derived from Fig. 3b), A3H-II/GFP (dark blue, input: 52 factors derived from Fig. 3b) and proteins shared between both A3H-I and A3H-II/GFP (“common”, medium blue, input: 121 factors derived from Fig. 3b) (LFC > 1, p-value < 0.01). **(c)** Polyclonal RKO-DOX-TID-A3H-I/II/GFP cells were treated with different concentrations of DOX for two days to achieve similar protein levels. Subsequently, cells were treated with EPOX for 5 h., during the last 15 min. of which, exogenous biotin was added to the culture media. Biotinylated proteins were purified, and their interaction with endogenous UBR5 and HUWE1 detected by WB. Relative abundance of UBR5/MYC (TID) or HUWE1/MYC (TID) was quantified. Densitometry values are listed.

**Figure S4.**
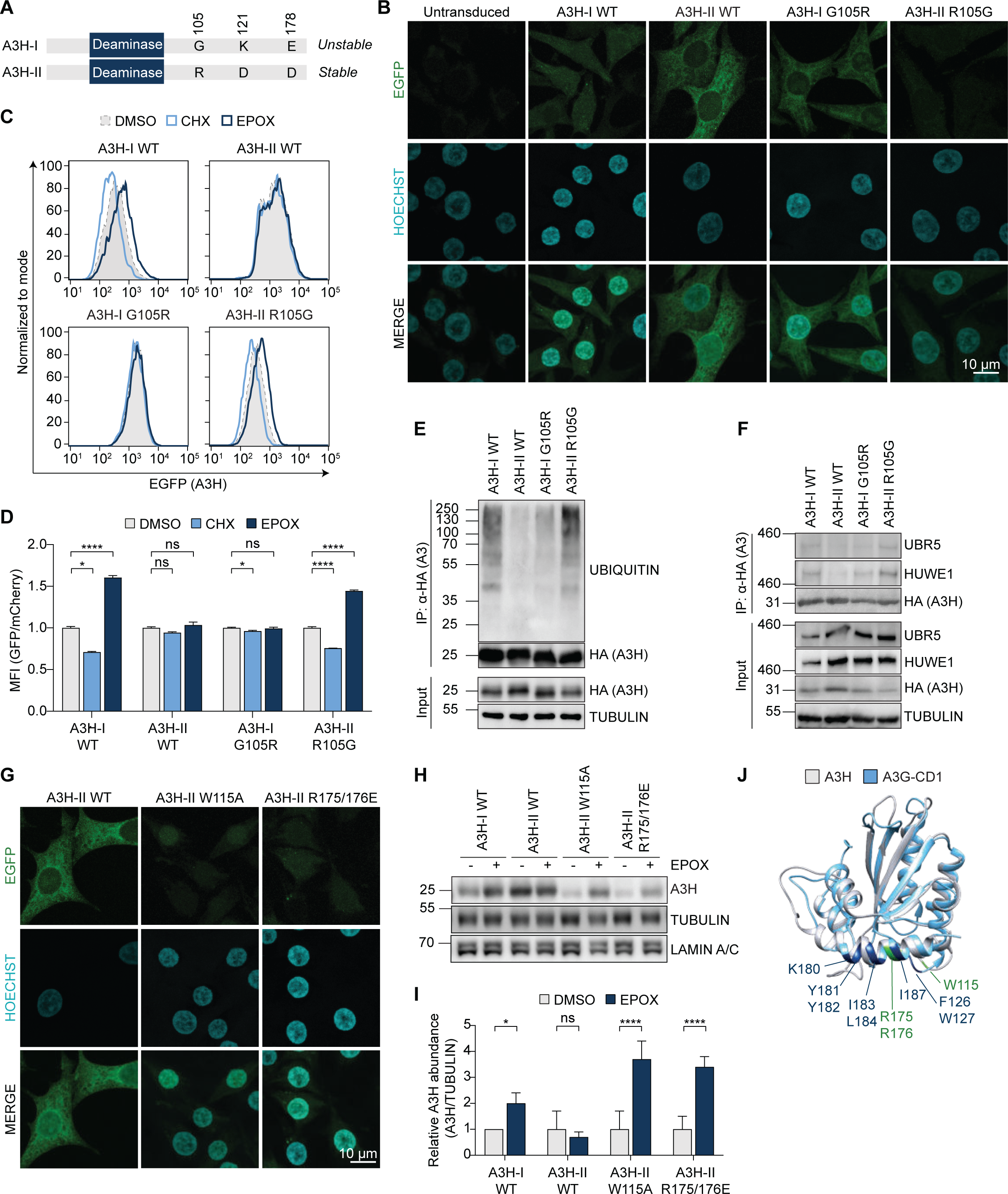
RNA binding protects A3 from E3 ligase binding and ubiquitination, thus promoting stability in cells. **(a)** Schematic overview of the three differential amino acid positions in A3H-I and A3H-II. **(b)** Confocal microscopy of RKO-mCherry-P2A-EGFP-A3H-I/II/I-G105R/II-R105G cells. Maximum intensity projections of 11 slices are displayed (scale bar = 10 µm). **(c-d)** RKO-mCherry-P2A-EGFP-A3H cells expressing the indicated EGFP-tagged A3H variants were treated for 5 h. with EPOX or CHX, followed by measurement of the mean fluorescence intensity (MFI) of EGFP-A3H by flow cytometry. **(d)** Quantification of (c) (means and SD, one-way ANOVA, **** p < 0.0001, * p < 0.05 n = 3). **(e-f)** Different amounts of the indicated 3xHA-A3H plasmids were transiently transfected in HEK-293T cells to achieve similar steady-state A3 protein levels, followed by 5 h. of EPOX treatment and immunoprecipitation of 3xHA-tagged proteins from the cell lysates. Their **(e)** ubiquitination or **(f)** interaction with UBR5 and HUWE1 was analyzed by WB. **(g)** Confocal microscopy of RKO-mCherry-P2A-EGFP-A3H cells expressing the indicated EGFP-tagged A3H variants. Maximum intensity projections of 11 slices are displayed (scale bar = 10 µm). **(h)** Whole cell extract samples corresponding to Fig. 4c. **(i)** Quantification of (h) (multiple unpaired t-tests, **** p < 0.0001, * p < 0.05; n = 3). **(j)** Structural alignment of A3H-II (PDB:6B0B) and A3G-CD1 (PDB: 5K81). Relevant amino acid residues mutated in the various RNA-binding mutants are colored in green (A3H-II) or dark blue (A3G).

**Figure S5.**
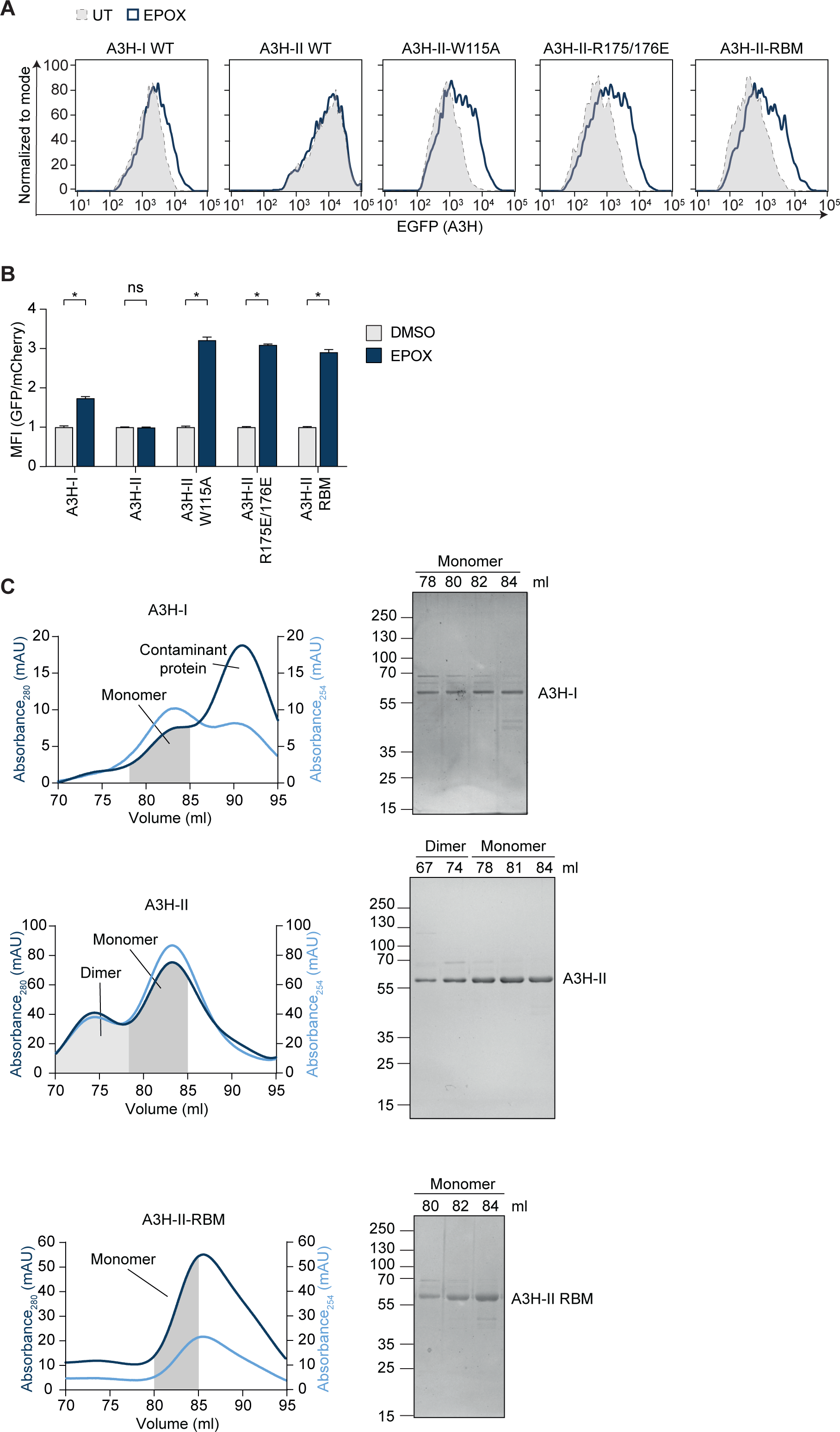
RNA binding by APOBEC3 proteins prevents their recognition and ubiquitination by E3 ligases *in vitro*. **(a-b)** HEK-293T cells were transiently transfected with EGFP-tagged A3H-I/II or A3H-II-RBM. After 48 h., cells were treated with EPOX for 5 h., followed by **(a)** measurement of EGFP-A3H fluorescence by flow cytometry, and **b)** followed by subsequent quantification (multiple unpaired t-tests, * p < 0.05, n = 3). **(c)** SEC profiles of 10x-His-MBP-A3H-I, 10x-His-MBP-A3H-II and 10x-His-MBP-A3H-II-RBM. Dark blue lines depict the absorbance at 280 nm representing eluted protein, light blue lines at 254 nm representing nucleic acid bound to the protein. Monomeric or dimeric fractions were pooled separately and analyzed by Coomassie stained SDS PAGE (right insets).

**Figure S6.**
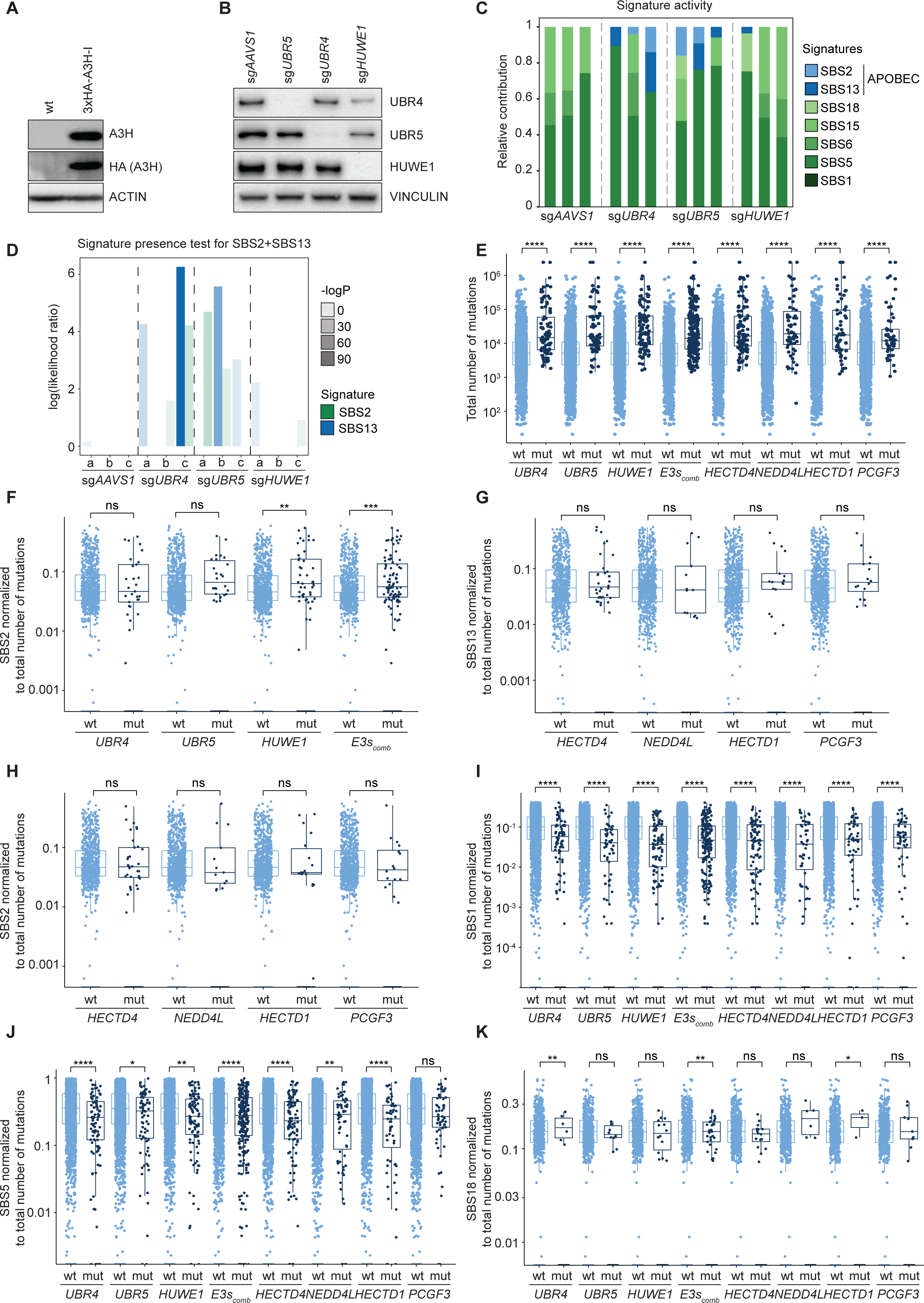
E3 ligase loss or mutation increases APOBEC signature mutations. **(a)** Cell lysates from a *UNG2*^-/-^ RKO cell line expressing DOX-inducible Cas9 and mCherry-P2A-3xHA-A3H-I were analyzed by WB for transgene expression. **(b)** These cells we transduced with sgRNAs targeting *UBR4*, *UBR5*, or *HUWE1*, and subsequently sorted for sgRNA-positive cells. Gene editing was induced with DOX for up to four cell doublings, after which the protein levels of the targeted E3 ligases were analyzed by WB. **(c)** Best subset signature refitting of mutREAD data, averaged per genotype, using signatures related to overactivity of APOBEC family enzymes and signatures commonly active in colon carcinoma, the parent tumor type for the model cell line RKO. For the purpose of comparing signature activity across genotypes, samples were matched by doubling time. Each bar represents the signature refitting results, scaled to 1 to show the relative signature contribution. **(d)** Signature presence test for the indicated signatures. Genotypes (n = 3, a-c), are divided by vertical dashed lines. The y-axis shows the log of the likelihood ratio, representing the maximum likelihood of the data, given a refit model including the signature of interest over the maximum likelihood of the data, given a refit model excluding the signature of interest. A log(likelihood ratio) greater than 0 indicates a significant activity of the signature of interest. The translucence of the bars (-log(p)) indicates the level of significance of the likelihood ratio test compared to the null hypothesis, which assumes that the mutational profile can be reasonably reconstructed without the signature of interest. **(e-k)** Cancers from TCGA/ICGC were grouped, based on whether the indicated E3 ligase genes were wild-type (wt) or mutated (mut), and the **(e)** total number of mutations per group plotted. **(f-k)** Subsequently, mutational signatures were normalized to the total number of mutations in each sample, and the levels of indicated signatures compared between two groups. “E3s_comb_” compares all samples, in which *UBR4*, *UBR5* and *HUWE1* are either all wild-type (wt), or at least one of the E3 ligase genes was mutated (mut). Data represent Wilcox rank sum test between wt and mut of each genotype, ns: p > 0.05, *: p ≤ 0.05, **: p ≤ 0.01, ***: p ≤ 0.001, ****: p ≤ 0.0001, n = 2703 cancer genome samples).

## References

1. Refsland, E.W., and Harris, R.S. (2013). The APOBEC3 family of retroelement restriction factors. Curr Top Microbiol Immunol 371, 1–27. 10.1007/978-3-642-37765-5_1.

2. Sheehy, A.M., Gaddis, N.C., Choi, J.D., and Malim, M.H. (2002). Isolation of a human gene that inhibits HIV-1 infection and is suppressed by the viral Vif protein. Nature 418, 646–650. 10.1038/nature00939.

3. Hultquist, J.F., Lengyel, J.A., Refsland, E.W., LaRue, R.S., Lackey, L., Brown, W.L., and Harris, R.S. (2011). Human and rhesus APOBEC3D, APOBEC3F, APOBEC3G, and APOBEC3H demonstrate a conserved capacity to restrict Vif-deficient HIV-1. J Virol 85, 11220–11234. 10.1128/JVI.05238-11.

4. Harris, R.S., Bishop, K.N., Sheehy, A.M., Craig, H.M., Petersen-Mahrt, S.K., Watt, I.N., Neuberger, M.S., and Malim, M.H. (2003). DNA deamination mediates innate immunity to retroviral infection. Cell 113, 803–809. 10.1016/s0092-8674(03)00423-9.

5. Lecossier, D., Bouchonnet, F., Clavel, F., and Hance, A.J. (2003). Hypermutation of HIV-1 DNA in the absence of the Vif protein. Science 300, 1112. 10.1126/science.1083338.

6. Mangeat, B., Turelli, P., Caron, G., Friedli, M., Perrin, L., and Trono, D. (2003). Broad antiretroviral defence by human APOBEC3G through lethal editing of nascent reverse transcripts. Nature 424, 99–103. 10.1038/nature01709.

7. Zhang, H., Yang, B., Pomerantz, R.J., Zhang, C., Arunachalam, S.C., and Gao, L. (2003). The cytidine deaminase CEM15 induces hypermutation in newly synthesized HIV-1 DNA. Nature 424, 94–98. 10.1038/nature01707.

8. Rogozin, I.B., Iyer, L.M., Liang, L., Glazko, G.V., Liston, V.G., Pavlov, Y.I., Aravind, L., and Pancer, Z. (2007). Evolution and diversification of lamprey antigen receptors: evidence for involvement of an AID-APOBEC family cytosine deaminase. Nat Immunol 8, 647–656. 10.1038/ni1463.

9. Conticello, S.G. (2008). The AID/APOBEC family of nucleic acid mutators. Genome Biol 9, 229. 10.1186/gb-2008-9-6-229.

10. Wang, X., Abudu, A., Son, S., Dang, Y., Venta, P.J., and Zheng, Y.-H. (2011). Analysis of human APOBEC3H haplotypes and anti-human immunodeficiency virus type 1 activity. J Virol 85, 3142–3152. 10.1128/JVI.02049-10.

11. Harari, A., Ooms, M., Mulder, L.C.F., and Simon, V. (2009). Polymorphisms and splice variants influence the antiretroviral activity of human APOBEC3H. J Virol 83, 295–303. 10.1128/JVI.01665-08.

12. OhAinle, M., Kerns, J.A., Li, M.M.H., Malik, H.S., and Emerman, M. (2008). Antiretroelement activity of APOBEC3H was lost twice in recent human evolution. Cell Host Microbe 4, 249–259. 10.1016/j.chom.2008.07.005.

13. Ooms, M., Majdak, S., Seibert, C.W., Harari, A., and Simon, V. (2010). The Localization of APOBEC3H Variants in HIV-1 Virions Determines Their Antiviral Activity. J Virol 84, 7961–7969. 10.1128/JVI.00754-10.

14. Li, M.M.H., and Emerman, M. (2011). Polymorphism in human APOBEC3H affects a phenotype dominant for subcellular localization and antiviral activity. J Virol 85, 8197– 8207. 10.1128/JVI.00624-11.

15. DeWeerd, R.A., Németh, E., Póti, Á., Petryk, N., Chen, C.-L., Hyrien, O., Szüts, D., and Green, A.M. (2022). Prospectively defined patterns of APOBEC3A mutagenesis are prevalent in human cancers. Cell Reports 38, 110555. 10.1016/j.celrep.2022.110555.

16. Law, E.K., Levin-Klein, R., Jarvis, M.C., Kim, H., Argyris, P.P., Carpenter, M.A., Starrett, G.J., Temiz, N.A., Larson, L.K., Durfee, C., et al. (2020). APOBEC3A catalyzes mutation and drives carcinogenesis in vivo. J Exp Med 217, e20200261. 10.1084/jem.20200261.

17. Petljak, M., Dananberg, A., Chu, K., Bergstrom, E.N., Striepen, J., von Morgen, P., Chen, Y., Shah, H., Sale, J.E., Alexandrov, L.B., et al. (2022). Mechanisms of APOBEC3 mutagenesis in human cancer cells. Nature 607, 799–807. 10.1038/s41586-022-04972-y.

18. Starrett, G.J., Luengas, E.M., McCann, J.L., Ebrahimi, D., Temiz, N.A., Love, R.P., Feng, Y., Adolph, M.B., Chelico, L., Law, E.K., et al. (2016). The DNA cytosine deaminase APOBEC3H haplotype I likely contributes to breast and lung cancer mutagenesis. Nat Commun 7, 12918. 10.1038/ncomms12918.

19. Henderson, S., and Fenton, T. (2015). APOBEC3 genes: retroviral restriction factors to cancer drivers. Trends Mol Med 21, 274–284. 10.1016/j.molmed.2015.02.007.

20. Jalili, P., Bowen, D., Langenbucher, A., Park, S., Aguirre, K., Corcoran, R.B., Fleischman, A.G., Lawrence, M.S., Zou, L., and Buisson, R. (2020). Quantification of ongoing APOBEC3A activity in tumor cells by monitoring RNA editing at hotspots. Nat Commun 11, 2971. 10.1038/s41467-020-16802-8.

21. Petljak, M., Alexandrov, L.B., Brammeld, J.S., Price, S., Wedge, D.C., Grossmann, S., Dawson, K.J., Ju, Y.S., Iorio, F., Tubio, J.M.C., et al. (2019). Characterizing Mutational Signatures in Human Cancer Cell Lines Reveals Episodic APOBEC Mutagenesis. Cell 176, 1282–1294.e20. 10.1016/j.cell.2019.02.012.

22. Nik-Zainal, S., Alexandrov, L.B., Wedge, D.C., Van Loo, P., Greenman, C.D., Raine, K., Jones, D., Hinton, J., Marshall, J., Stebbings, L.A., et al. (2012). Mutational processes molding the genomes of 21 breast cancers. Cell 149, 979–993. 10.1016/j.cell.2012.04.024.

23. Nik-Zainal, S., Davies, H., Staaf, J., Ramakrishna, M., Glodzik, D., Zou, X., Martincorena, I., Alexandrov, L.B., Martin, S., Wedge, D.C., et al. (2016). Landscape of somatic mutations in 560 breast cancer whole-genome sequences. Nature 534, 47–54. 10.1038/nature17676.

24. Alexandrov, L.B., Nik-Zainal, S., Wedge, D.C., Aparicio, S.A.J.R., Behjati, S., Biankin, A.V., Bignell, G.R., Bolli, N., Borg, A., Børresen-Dale, A.-L., et al. (2013). Signatures of mutational processes in human cancer. Nature 500, 415–421. 10.1038/nature12477.

25. Roberts, S.A., Lawrence, M.S., Klimczak, L.J., Grimm, S.A., Fargo, D., Stojanov, P., Kiezun, A., Kryukov, G.V., Carter, S.L., Saksena, G., et al. (2013). An APOBEC cytidine deaminase mutagenesis pattern is widespread in human cancers. Nat Genet 45, 970– 976. 10.1038/ng.2702.

26. de Bruin, E.C., McGranahan, N., Mitter, R., Salm, M., Wedge, D.C., Yates, L., Jamal-Hanjani, M., Shafi, S., Murugaesu, N., Rowan, A.J., et al. (2014). Spatial and temporal diversity in genomic instability processes defines lung cancer evolution. Science 346, 251–256. 10.1126/science.1253462.

27. Morganella, S., Alexandrov, L.B., Glodzik, D., Zou, X., Davies, H., Staaf, J., Sieuwerts, A.M., Brinkman, A.B., Martin, S., Ramakrishna, M., et al. (2016). The topography of mutational processes in breast cancer genomes. Nat Commun 7, 11383. 10.1038/ncomms11383.

28. Carpenter, M.A., Temiz, N.A., Ibrahim, M.A., Jarvis, M.C., Brown, M.R., Argyris, P.P., Brown, W.L., Starrett, G.J., Yee, D., and Harris, R.S. (2023). Mutational impact of APOBEC3A and APOBEC3B in a human cell line and comparisons to breast cancer. PLoS Genet 19, e1011043. 10.1371/journal.pgen.1011043.

29. Bergstrom, E.N., Luebeck, J., Petljak, M., Khandekar, A., Barnes, M., Zhang, T., Steele, C.D., Pillay, N., Landi, M.T., Bafna, V., et al. (2022). Mapping clustered mutations in cancer reveals APOBEC3 mutagenesis of ecDNA. Nature 602, 510–517. 10.1038/s41586-022-04398-6.

30. Alexandrov, L.B., Kim, J., Haradhvala, N.J., Huang, M.N., Tian Ng, A.W., Wu, Y., Boot, A., Covington, K.R., Gordenin, D.A., Bergstrom, E.N., et al. (2020). The repertoire of mutational signatures in human cancer. Nature 578, 94–101. 10.1038/s41586-020-1943-3.

31. Dananberg, A., Striepen, J., Rozowsky, J.S., and Petljak, M. (2024). APOBEC Mutagenesis in Cancer Development and Susceptibility. Cancers (Basel) 16, 374. 10.3390/cancers16020374.

32. Hix, M.A., Wong, L., Flath, B., Chelico, L., and Cisneros, G.A. (2020). Single-nucleotide polymorphism of the DNA cytosine deaminase APOBEC3H haplotype I leads to enzyme destabilization and correlates with lung cancer. NAR Cancer 2, zcaa023. 10.1093/narcan/zcaa023.

33. Sanchez, A., Ortega, P., Sakhtemani, R., Manjunath, L., Oh, S., Bournique, E., Becker, A., Kim, K., Durfee, C., Temiz, N.A., et al. (2024). Mesoscale DNA features impact APOBEC3A and APOBEC3B deaminase activity and shape tumor mutational landscapes. Nat Commun 15, 2370. 10.1038/s41467-024-45909-5.

34. Burns, M.B., Lackey, L., Carpenter, M.A., Rathore, A., Land, A.M., Leonard, B., Refsland, E.W., Kotandeniya, D., Tretyakova, N., Nikas, J.B., et al. (2013). APOBEC3B is an enzymatic source of mutation in breast cancer. Nature 494, 366–370. 10.1038/nature11881.

35. Burns, M.B., Temiz, N.A., and Harris, R.S. (2013). Evidence for APOBEC3B mutagenesis in multiple human cancers. Nat Genet 45, 977–983. 10.1038/ng.2701.

36. Apolonia, L., Schulz, R., Curk, T., Rocha, P., Swanson, C.M., Schaller, T., Ule, J., and Malim, M.H. (2015). Promiscuous RNA binding ensures effective encapsidation of APOBEC3 proteins by HIV-1. PLoS Pathog 11, e1004609. 10.1371/journal.ppat.1004609.

37. York, A., Kutluay, S.B., Errando, M., and Bieniasz, P.D. (2016). The RNA Binding Specificity of Human APOBEC3 Proteins Resembles That of HIV-1 Nucleocapsid. PLoS Pathog 12, e1005833. 10.1371/journal.ppat.1005833.

38. Dang, Y., Siew, L.M., Wang, X., Han, Y., Lampen, R., and Zheng, Y.-H. (2008). Human cytidine deaminase APOBEC3H restricts HIV-1 replication. J Biol Chem 283, 11606– 11614. 10.1074/jbc.M707586200.

39. Shaban, N.M., Shi, K., Lauer, K.V., Carpenter, M.A., Richards, C.M., Salamango, D., Wang, J., Lopresti, M.W., Banerjee, S., Levin-Klein, R., et al. (2018). The Antiviral and Cancer Genomic DNA Deaminase APOBEC3H Is Regulated by an RNA-Mediated Dimerization Mechanism. Mol Cell 69, 75–86.e9. 10.1016/j.molcel.2017.12.010.

40. Refsland, E.W., Hultquist, J.F., Luengas, E.M., Ikeda, T., Shaban, N.M., Law, E.K., Brown, W.L., Reilly, C., Emerman, M., and Harris, R.S. (2014). Natural polymorphisms in human APOBEC3H and HIV-1 Vif combine in primary T lymphocytes to affect viral G-to-A mutation levels and infectivity. PLoS Genet 10, e1004761. 10.1371/journal.pgen.1004761.

41. Lackey, L., Demorest, Z.L., Land, A.M., Hultquist, J.F., Brown, W.L., and Harris, R.S. (2012). APOBEC3B and AID have similar nuclear import mechanisms. J Mol Biol 419, 301–314. 10.1016/j.jmb.2012.03.011.

42. Salamango, D.J., Becker, J.T., McCann, J.L., Cheng, A.Z., Demir, Ö., Amaro, R.E., Brown, W.L., Shaban, N.M., and Harris, R.S. (2018). APOBEC3H Subcellular Localization Determinants Define Zipcode for Targeting HIV-1 for Restriction. Mol Cell Biol 38, e00356–18. 10.1128/MCB.00356-18.

43. Chesarino, N.M., and Emerman, M. (2020). Polymorphisms in Human APOBEC3H Differentially Regulate Ubiquitination and Antiviral Activity. Viruses 12, 378. 10.3390/v12040378.

44. de Almeida, M., Hinterndorfer, M., Brunner, H., Grishkovskaya, I., Singh, K., Schleiffer, A., Jude, J., Deswal, S., Kalis, R., Vunjak, M., et al. (2021). AKIRIN2 controls the nuclear import of proteasomes in vertebrates. Nature 599, 491–496. 10.1038/s41586-021-04035-8.

45. Vunjak, M., Schwartz, I., Cantoran García, A., Mastrovito, M., Hinterndorfer, M., de Almeida, M., Budroni, V., Wang, J., Froussios, K., Jude, J., et al. (2022). SPOP targets the immune transcription factor IRF1 for proteasomal degradation (Cell Biology) 10.1101/2022.10.10.511567.

46. Scinicariello, S., Soderholm, A., Schäfer, M., Shulkina, A., Schwartz, I., Hacker, K., Gogova, R., Kalis, R., Froussios, K., Budroni, V., et al. (2023). HUWE1 controls tristetraprolin proteasomal degradation by regulating its phosphorylation. Elife 12, e83159. 10.7554/eLife.83159.

47. Jang, G.M., Annan Sudarsan, A.K., Shayeganmehr, A., Prando Munhoz, E., Lao, R., Gaba, A., Granadillo Rodríguez, M., Love, R.P., Polacco, B.J., Zhou, Y., et al. (2024). Protein interaction map of APOBEC3 enzyme family reveals deamination-independent role in cellular function. Mol Cell Proteomics, 100755. 10.1016/j.mcpro.2024.100755.

48. Gallois-Montbrun, S., Kramer, B., Swanson, C.M., Byers, H., Lynham, S., Ward, M., and Malim, M.H. (2007). Antiviral protein APOBEC3G localizes to ribonucleoprotein complexes found in P bodies and stress granules. J Virol 81, 2165–2178. 10.1128/JVI.02287-06.

49. Izumi, T., Burdick, R., Shigemi, M., Plisov, S., Hu, W.-S., and Pathak, V.K. (2013). Mov10 and APOBEC3G localization to processing bodies is not required for virion incorporation and antiviral activity. J Virol 87, 11047–11062. 10.1128/JVI.02070-13.

50. Phalora, P.K., Sherer, N.M., Wolinsky, S.M., Swanson, C.M., and Malim, M.H. (2012). HIV-1 replication and APOBEC3 antiviral activity are not regulated by P bodies. J Virol 86, 11712–11724. 10.1128/JVI.00595-12.

51. Kozak, S.L., Marin, M., Rose, K.M., Bystrom, C., and Kabat, D. (2006). The anti-HIV-1 editing enzyme APOBEC3G binds HIV-1 RNA and messenger RNAs that shuttle between polysomes and stress granules. J Biol Chem 281, 29105–29119. 10.1074/jbc.M601901200.

52. Ito, F., Yang, H., Xiao, X., Li, S.-X., Wolfe, A., Zirkle, B., Arutiunian, V., and Chen, X.S. (2018). Understanding the Structure, Multimerization, Subcellular Localization and mC Selectivity of a Genomic Mutator and Anti-HIV Factor APOBEC3H. Sci Rep 8, 3763. 10.1038/s41598-018-21955-0.

53. Zhen, A., Du, J., Zhou, X., Xiong, Y., and Yu, X.-F. (2012). Reduced APOBEC3H variant anti-viral activities are associated with altered RNA binding activities. PLoS One 7, e38771. 10.1371/journal.pone.0038771.

54. Polevoda, B., Joseph, R., Friedman, A.E., Bennett, R.P., Greiner, R., De Zoysa, T., Stewart, R.A., and Smith, H.C. (2017). DNA mutagenic activity and capacity for HIV-1 restriction of the cytidine deaminase APOBEC3G depend on whether DNA or RNA binds to tyrosine 315. Journal of Biological Chemistry 292, 8642–8656. 10.1074/jbc.M116.767889.

55. Matsuoka, T., Nagae, T., Ode, H., Awazu, H., Kurosawa, T., Hamano, A., Matsuoka, K., Hachiya, A., Imahashi, M., Yokomaku, Y., et al. (2018). Structural basis of chimpanzee APOBEC3H dimerization stabilized by double-stranded RNA. Nucleic Acids Research 46, 10368–10379. 10.1093/nar/gky676.

56. Bohn, J.A., Thummar, K., York, A., Raymond, A., Brown, W.C., Bieniasz, P.D., Hatziioannou, T., and Smith, J.L. (2017). APOBEC3H structure reveals an unusual mechanism of interaction with duplex RNA. Nat Commun 8, 1021. 10.1038/s41467-017-01309-6.

57. Yau, R.G., Doerner, K., Castellanos, E.R., Haakonsen, D.L., Werner, A., Wang, N., Yang, X.W., Martinez-Martin, N., Matsumoto, M.L., Dixit, V.M., et al. (2017). Assembly and Function of Heterotypic Ubiquitin Chains in Cell-Cycle and Protein Quality Control. Cell 171, 918–933.e20. 10.1016/j.cell.2017.09.040.

58. Perner, J., Abbas, S., Nowicki-Osuch, K., Devonshire, G., Eldridge, M.D., Tavaré, S., and Fitzgerald, R.C. (2020). The mutREAD method detects mutational signatures from low quantities of cancer DNA. Nat Commun 11, 3166. 10.1038/s41467-020-16974-3.

59. The ICGC/TCGA Pan-Cancer Analysis of Whole Genomes Consortium, Aaltonen, L.A., Abascal, F., Abeshouse, A., Aburatani, H., Adams, D.J., Agrawal, N., Ahn, K.S., Ahn, S.-M., Aikata, H., et al. (2020). Pan-cancer analysis of whole genomes. Nature 578, 82–93. 10.1038/s41586-020-1969-6.

60. Butler, K., and Banday, A.R. (2023). APOBEC3-mediated mutagenesis in cancer: causes, clinical significance and therapeutic potential. J Hematol Oncol 16, 31. 10.1186/s13045-023-01425-5.

61. Durfee, C., Temiz, N.A., Levin-Klein, R., Argyris, P.P., Alsøe, L., Carracedo, S., Alonso de la Vega, A., Proehl, J., Holzhauer, A.M., Seeman, Z.J., et al. (2023). Human APOBEC3B promotes tumor development in vivo including signature mutations and metastases. Cell Rep Med 4, 101211. 10.1016/j.xcrm.2023.101211.

62. McCann, J.L., Cristini, A., Law, E.K., Lee, S.Y., Tellier, M., Carpenter, M.A., Beghè, C., Kim, J.J., Sanchez, A., Jarvis, M.C., et al. (2023). APOBEC3B regulates R-loops and promotes transcription-associated mutagenesis in cancer. Nat Genet 55, 1721–1734. 10.1038/s41588-023-01504-w.

63. Middlebrooks, C.D., Banday, A.R., Matsuda, K., Udquim, K.-I., Onabajo, O.O., Paquin, A., Figueroa, J.D., Zhu, B., Koutros, S., Kubo, M., et al. (2016). Association of germline variants in the APOBEC3 region with cancer risk and enrichment with APOBEC-signature mutations in tumors. Nat Genet 48, 1330–1338. 10.1038/ng.3670.

64. Law, E.K., Sieuwerts, A.M., LaPara, K., Leonard, B., Starrett, G.J., Molan, A.M., Temiz, N.A., Vogel, R.I., Meijer-van Gelder, M.E., Sweep, F.C.G.J., et al. (2016). The DNA cytosine deaminase APOBEC3B promotes tamoxifen resistance in ER-positive breast cancer. Sci Adv 2, e1601737. 10.1126/sciadv.1601737.

65. Stopak, K.S., Chiu, Y.-L., Kropp, J., Grant, R.M., and Greene, W.C. (2007). Distinct patterns of cytokine regulation of APOBEC3G expression and activity in primary lymphocytes, macrophages, and dendritic cells. J Biol Chem 282, 3539–3546. 10.1074/jbc.M610138200.

66. Kreisberg, J.F., Yonemoto, W., and Greene, W.C. (2006). Endogenous factors enhance HIV infection of tissue naive CD4 T cells by stimulating high molecular mass APOBEC3G complex formation. J Exp Med 203, 865–870. 10.1084/jem.20051856.

67. Soros, V.B., Yonemoto, W., and Greene, W.C. (2007). Newly synthesized APOBEC3G is incorporated into HIV virions, inhibited by HIV RNA, and subsequently activated by RNase H. PLoS Pathog 3, e15. 10.1371/journal.ppat.0030015.

68. Friew, Y.N., Boyko, V., Hu, W.-S., and Pathak, V.K. (2009). Intracellular interactions between APOBEC3G, RNA, and HIV-1 Gag: APOBEC3G multimerization is dependent on its association with RNA. Retrovirology 6, 56. 10.1186/1742-4690-6-56.

69. Khan, M.A., Goila-Gaur, R., Opi, S., Miyagi, E., Takeuchi, H., Kao, S., and Strebel, K. (2007). Analysis of the contribution of cellular and viral RNA to the packaging of APOBEC3G into HIV-1 virions. Retrovirology 4, 48. 10.1186/1742-4690-4-48.

70. Wittkopp, C.J., Adolph, M.B., Wu, L.I., Chelico, L., and Emerman, M. (2016). A Single Nucleotide Polymorphism in Human APOBEC3C Enhances Restriction of Lentiviruses. PLoS Pathog 12, e1005865. 10.1371/journal.ppat.1005865.

71. Monda, J.K., Ge, X., Hunkeler, M., Donovan, K.A., Ma, M.W., Jin, C.Y., Leonard, M., Fischer, E.S., and Bennett, E.J. (2023). HAPSTR1 localizes HUWE1 to the nucleus to limit stress signaling pathways. Cell Reports 42. 10.1016/j.celrep.2023.112496.

72. Xu, Y., Anderson, D.E., and Ye, Y. (2016). The HECT domain ubiquitin ligase HUWE1 targets unassembled soluble proteins for degradation. Cell Discov 2, 1–16. 10.1038/celldisc.2016.40.

73. Tripathi, S., Pohl, M.O., Zhou, Y., Rodriguez-Frandsen, A., Wang, G., Stein, D.A., Moulton, H.M., DeJesus, P., Che, J., Mulder, L.C.F., et al. (2015). Meta-and Orthogonal Integration of Influenza “OMICs” Data Defines a Role for UBR4 in Virus Budding. Cell Host & Microbe 18, 723–735. 10.1016/j.chom.2015.11.002.

74. Kim, S.T., Lee, Y.J., Tasaki, T., Hwang, J., Kang, M.J., Yi, E.C., Kim, B.Y., and Kwon, Y.T. (2018). The N-recognin UBR4 of the N-end rule pathway is required for neurogenesis and homeostasis of cell surface proteins. PLOS ONE 13, e0202260. 10.1371/journal.pone.0202260.

75. Chen, L.J., Xu, W.M., Yang, M., Wang, K., Chen, Y., Huang, X.J., and Ma, Q.H. (2016). HUWE1 plays important role in mouse preimplantation embryo development and the dysregulation is associated with poor embryo development in humans. Sci Rep 6, 37928. 10.1038/srep37928.

76. Shearer, R.F., Frikstad, K.-A.M., McKenna, J., McCloy, R.A., Deng, N., Burgess, A., Stokke, T., Patzke, S., and Saunders, D.N. (2018). The E3 ubiquitin ligase UBR5 regulates centriolar satellite stability and primary cilia. MBoC 29, 1542–1554. 10.1091/mbc.E17-04-0248.

77. Sanchez, A., De Vivo, A., Uprety, N., Kim, J., Stevens, S.M., and Kee, Y. (2016). BMI1– UBR5 axis regulates transcriptional repression at damaged chromatin. Proceedings of the National Academy of Sciences 113, 11243–11248. 10.1073/pnas.1610735113.

78. Xiang, G., Wang, S., Chen, L., Song, M., Song, X., Wang, H., Zhou, P., Ma, X., and Yu, J. (2022). UBR5 targets tumor suppressor CDC73 proteolytically to promote aggressive breast cancer. Cell Death Dis 13, 1–14. 10.1038/s41419-022-04914-6.

79. Jang, G.M., Sudarsan, A.K.A., Shayeganmehr, A., Munhoz, E.P., Lao, R., Gaba, A., Rodríguez, M.G., Love, R.P., Polacco, B.J., Zhou, Y., et al. (2024). Protein interaction map of APOBEC3 enzyme family reveals deamination-independent role in cellular function. bioRxiv, 2024.02.06.579137. 10.1101/2024.02.06.579137.

80. Esnault, C., Millet, J., Schwartz, O., and Heidmann, T. (2006). Dual inhibitory effects of APOBEC family proteins on retrotransposition of mammalian endogenous retroviruses. Nucleic Acids Res 34, 1522–1531. 10.1093/nar/gkl054.

81. Nakaya, Y., Stavrou, S., Blouch, K., Tattersall, P., and Ross, S.R. (2016). In Vivo Examination of Mouse APOBEC3-and Human APOBEC3A-and APOBEC3G-Mediated Restriction of Parvovirus and Herpesvirus Infection in Mouse Models. J Virol 90, 8005– 8012. 10.1128/JVI.00973-16.

82. Moraes, S.N., Becker, J.T., Moghadasi, S.A., Shaban, N.M., Auerbach, A.A., Cheng, A.Z., and Harris, R.S. (2022). Evidence linking APOBEC3B genesis and evolution of innate immune antagonism by gamma-herpesvirus ribonucleotide reductases. Elife 11, e83893. 10.7554/eLife.83893.

83. Cheng, A.Z., Yockteng-Melgar, J., Jarvis, M.C., Malik-Soni, N., Borozan, I., Carpenter, M.A., McCann, J.L., Ebrahimi, D., Shaban, N.M., Marcon, E., et al. (2019). Epstein-Barr virus BORF2 inhibits cellular APOBEC3B to preserve viral genome integrity. Nat Microbiol 4, 78–88. 10.1038/s41564-018-0284-6.

84. Zhang, Z., Gu, Q., de Manuel Montero, M., Bravo, I.G., Marques-Bonet, T., Häussinger, D., and Münk, C. (2017). Stably expressed APOBEC3H forms a barrier for cross-species transmission of simian immunodeficiency virus of chimpanzee to humans. PLoS Pathog 13, e1006746. 10.1371/journal.ppat.1006746.

85. Chen, Y., Hu, J., Cai, X., Huang, Y., Zhou, X., Tu, Z., Hu, J., Tavis, J.E., Tang, N., Huang, A., et al. (2018). APOBEC3B edits HBV DNA and inhibits HBV replication during reverse transcription. Antiviral Res 149, 16–25. 10.1016/j.antiviral.2017.11.006.

86. Bandarra, S., Miyagi, E., Ribeiro, A.C., Gonçalves, J., Strebel, K., and Barahona, I. (2021). APOBEC3B Potently Restricts HIV-2 but Not HIV-1 in a Vif-Dependent Manner. J Virol 95, e0117021. 10.1128/JVI.01170-21.

87. Mark, K.G., Kolla, S., Aguirre, J.D., Garshott, D.M., Schmitt, S., Haakonsen, D.L., Xu, C., Kater, L., Kempf, G., Martínez-González, B., et al. (2023). Orphan quality control shapes network dynamics and gene expression. Cell 186, 3460–3475.e23. 10.1016/j.cell.2023.06.015.

88. Tsai, J.M., Aguirre, J.D., Li, Y.-D., Brown, J., Focht, V., Kater, L., Kempf, G., Sandoval, B., Schmitt, S., Rutter, J.C., et al. (2023). UBR5 forms ligand-dependent complexes on chromatin to regulate nuclear hormone receptor stability. Mol Cell 83, 2753–2767.e10. 10.1016/j.molcel.2023.06.028.

89. Kaisari, S., Miniowitz-Shemtov, S., Sitry-Shevah, D., Shomer, P., Kozlov, G., Gehring, K., and Hershko, A. (2022). Role of ubiquitin-protein ligase UBR5 in the disassembly of mitotic checkpoint complexes. Proc Natl Acad Sci U S A 119, e2121478119. 10.1073/pnas.2121478119.

90. Hehl, L.A., Horn-Ghetko, D., Prabu, J.R., Vollrath, R., Vu, D.T., Pérez Berrocal, D.A., Mulder, M.P.C., van der Heden van Noort, G.J., and Schulman, B.A. (2024). Structural snapshots along K48-linked ubiquitin chain formation by the HECT E3 UBR5. Nat Chem Biol 20, 190–200. 10.1038/s41589-023-01414-2.

91. Hodáková, Z., Grishkovskaya, I., Brunner, H.L., Bolhuis, D.L., Belačić, K., Schleiffer, A., Kotisch, H., Brown, N.G., and Haselbach, D. (2023). Cryo-EM structure of the chain-elongating E3 ubiquitin ligase UBR5. EMBO J 42, e113348. 10.15252/embj.2022113348.

92. Wang, F., He, Q., Zhan, W., Yu, Z., Finkin-Groner, E., Ma, X., Lin, G., and Li, H. (2023). Structure of the human UBR5 E3 ubiquitin ligase. Structure 31, 541–552.e4. 10.1016/j.str.2023.03.010.

93. Heidelberger, J.B., Voigt, A., Borisova, M.E., Petrosino, G., Ruf, S., Wagner, S.A., and Beli, P. (2018). Proteomic profiling of VCP substrates links VCP to K6-linked ubiquitylation and c-Myc function. EMBO Rep 19, e44754. 10.15252/embr.201744754.

94. Hong, J.H., Kaustov, L., Coyaud, E., Srikumar, T., Wan, J., Arrowsmith, C., and Raught, B. (2015). KCMF1 (potassium channel modulatory factor 1) Links RAD6 to UBR4 (ubiquitin N-recognin domain-containing E3 ligase 4) and lysosome-mediated degradation. Mol Cell Proteomics 14, 674–685. 10.1074/mcp.M114.042168.

95. Haakonsen, D.L., Heider, M., Ingersoll, A.J., Vodehnal, K., Witus, S.R., Uenaka, T., Wernig, M., and Rapé, M. (2024). Stress response silencing by an E3 ligase mutated in neurodegeneration. Nature 626, 874–880. 10.1038/s41586-023-06985-7.

96. Leto, D.E., Morgens, D.W., Zhang, L., Walczak, C.P., Elias, J.E., Bassik, M.C., and Kopito, R.R. (2019). Genome-wide CRISPR Analysis Identifies Substrate-Specific Conjugation Modules in ER-Associated Degradation. Mol Cell 73, 377–389.e11. 10.1016/j.molcel.2018.11.015.

97. Cassidy, K.B., Bang, S., Kurokawa, M., and Gerber, S.A. (2020). Direct regulation of Chk1 protein stability by E3 ubiquitin ligase HUWE1. FEBS J 287, 1985–1999. 10.1111/febs.15132.

98. Kunz, V., Bommert, K.S., Kruk, J., Schwinning, D., Chatterjee, M., Stühmer, T., Bargou, R., and Bommert, K. (2020). Targeting of the E3 ubiquitin-protein ligase HUWE1 impairs DNA repair capacity and tumor growth in preclinical multiple myeloma models. Sci Rep 10, 18419. 10.1038/s41598-020-75499-3.

99. Thompson, J.W., Nagel, J., Hoving, S., Gerrits, B., Bauer, A., Thomas, J.R., Kirschner, M.W., Schirle, M., and Luchansky, S.J. (2014). Quantitative Lys-ɛ-Gly-Gly (diGly) proteomics coupled with inducible RNAi reveals ubiquitin-mediated proteolysis of DNA damage-inducible transcript 4 (DDIT4) by the E3 ligase HUWE1. J Biol Chem 289, 28942–28955. 10.1074/jbc.M114.573352.

100. D’Arca, D., Zhao, X., Xu, W., Ramirez-Martinez, N.C., Iavarone, A., and Lasorella, A. (2010). Huwe1 ubiquitin ligase is essential to synchronize neuronal and glial differentiation in the developing cerebellum. Proc Natl Acad Sci U S A 107, 5875–5880. 10.1073/pnas.0912874107.

101. Poulsen, E.G., Steinhauer, C., Lees, M., Lauridsen, A.-M., Ellgaard, L., and Hartmann-Petersen, R. (2012). HUWE1 and TRIP12 collaborate in degradation of ubiquitin-fusion proteins and misframed ubiquitin. PLoS One 7, e50548. 10.1371/journal.pone.0050548.

102. Hegazi, S., Cheng, A.H., Krupp, J.J., Tasaki, T., Liu, J., Szulc, D.A., Ling, H.H., Rios Garcia, J., Seecharran, S., Basiri, T., et al. (2022). UBR4/POE facilitates secretory trafficking to maintain circadian clock synchrony. Nat Commun 13, 1594. 10.1038/s41467-022-29244-1.

103. Hunt, L.C., Stover, J., Haugen, B., Shaw, T.I., Li, Y., Pagala, V.R., Finkelstein, D., Barton, E.R., Fan, Y., Labelle, M., et al. (2019). A Key Role for the Ubiquitin Ligase UBR4 in Myofiber Hypertrophy in Drosophila and Mice. Cell Rep 28, 1268–1281.e6. 10.1016/j.celrep.2019.06.094.

104. Jeong, D.E., Lee, H.S., Ku, B., Kim, C.-H., Kim, S.J., and Shin, H.-C. (2023). Insights into the recognition mechanism in the UBR box of UBR4 for its specific substrates. Commun Biol 6, 1214. 10.1038/s42003-023-05602-7.

105. Rinschen, M.M., Bharill, P., Wu, X., Kohli, P., Reinert, M.J., Kretz, O., Saez, I., Schermer, B., Höhne, M., Bartram, M.P., et al. (2016). The ubiquitin ligase Ubr4 controls stability of podocin/MEC-2 supercomplexes. Hum Mol Genet 25, 1328–1344. 10.1093/hmg/ddw016.

106. Hunkeler, M., Jin, C.Y., Ma, M.W., Monda, J.K., Overwijn, D., Bennett, E.J., and Fischer, E.S. (2021). Solenoid architecture of HUWE1 contributes to ligase activity and substrate recognition. Mol Cell 81, 3468–3480.e7. 10.1016/j.molcel.2021.06.032.

107. Grabarczyk, D.B., Petrova, O.A., Deszcz, L., Kurzbauer, R., Murphy, P., Ahel, J., Vogel, A., Gogova, R., Faas, V., Kordic, D., et al. (2021). HUWE1 employs a giant substrate-binding ring to feed and regulate its HECT E3 domain. Nat Chem Biol 17, 1084–1092. 10.1038/s41589-021-00831-5.

108. Ohtake, F., Tsuchiya, H., Saeki, Y., and Tanaka, K. (2018). K63 ubiquitylation triggers proteasomal degradation by seeding branched ubiquitin chains. Proc Natl Acad Sci USA 115, E1401–E1408. 10.1073/pnas.1716673115.

109. Manasanch, E.E., and Orlowski, R.Z. (2017). Proteasome Inhibitors in Cancer Therapy. Nat Rev Clin Oncol 14, 417–433. 10.1038/nrclinonc.2016.206.

110. Venkatesan, S., Rosenthal, R., Kanu, N., McGranahan, N., Bartek, J., Quezada, S.A., Hare, J., Harris, R.S., and Swanton, C. (2018). Perspective: APOBEC mutagenesis in drug resistance and immune escape in HIV and cancer evolution. Ann Oncol 29, 563–572. 10.1093/annonc/mdy003.

111. Coxon, M., Dennis, M.A., Dananberg, A., Collins, C.D., Wilson, H.E., Meekma, J., Savenkova, M.I., Ng, D., Osbron, C.A., Mertz, T.M., et al. (2023). An impaired ubiquitin-proteasome system increases APOBEC3A abundance. NAR Cancer 5, zcad058. 10.1093/narcan/zcad058.

112. Shlyakhtenko, L.S., Lushnikov, A.J., Li, M., Harris, R.S., and Lyubchenko, Y.L. (2014). Interaction of APOBEC3A with DNA assessed by atomic force microscopy. PLoS One 9, e99354. 10.1371/journal.pone.0099354.

113. Chen, X.S. (2021). Insights into the Structures and Multimeric Status of APOBEC Proteins Involved in Viral Restriction and Other Cellular Functions. Viruses 13, 497. 10.3390/v13030497.

114. Bohn, M.-F., Shandilya, S.M.D., Silvas, T.V., Nalivaika, E.A., Kouno, T., Kelch, B.A., Ryder, S.P., Kurt-Yilmaz, N., Somasundaran, M., and Schiffer, C.A. (2015). The ssDNA Mutator APOBEC3A Is Regulated by Cooperative Dimerization. Structure 23, 903–911. 10.1016/j.str.2015.03.016.

115. Huthoff, H., Autore, F., Gallois-Montbrun, S., Fraternali, F., and Malim, M.H. (2009). RNA-Dependent Oligomerization of APOBEC3G Is Required for Restriction of HIV-1. PLoS Pathog 5, e1000330. 10.1371/journal.ppat.1000330.

116. Aydin, H., Taylor, M.W., and Lee, J.E. (2014). Structure-guided analysis of the human APOBEC3-HIV restrictome. Structure 22, 668–684. 10.1016/j.str.2014.02.011.

117. Michlits, G., Jude, J., Hinterndorfer, M., de Almeida, M., Vainorius, G., Hubmann, M., Neumann, T., Schleiffer, A., Burkard, T.R., Fellner, M., et al. (2020). Multilayered VBC score predicts sgRNAs that efficiently generate loss-of-function alleles. Nat Methods 17, 708–716. 10.1038/s41592-020-0850-8.

118. Schwartz, I., Vunjak, M., Budroni, V., Cantoran García, A., Mastrovito, M., Soderholm, A., Hinterndorfer, M., de Almeida, M., Hacker, K., Wang, J., et al. (2023). SPOP targets the immune transcription factor IRF1 for proteasomal degradation. Elife 12, e89951. 10.7554/eLife.89951.

119. Hatziioannou, T., Cowan, S., and Bieniasz, P.D. (2004). Capsid-dependent and - independent postentry restriction of primate lentivirus tropism in rodent cells. J Virol 78, 1006–1011. 10.1128/jvi.78.2.1006-1011.2004.

120. Yee, J.K., Friedmann, T., and Burns, J.C. (1994). Generation of high-titer pseudotyped retroviral vectors with very broad host range. Methods Cell Biol 43 Pt A, 99–112. 10.1016/s0091-679x(08)60600-7.

121. Li, W., Xu, H., Xiao, T., Cong, L., Love, M.I., Zhang, F., Irizarry, R.A., Liu, J.S., Brown, M., and Liu, X.S. (2014). MAGeCK enables robust identification of essential genes from genome-scale CRISPR/Cas9 knockout screens. Genome Biol 15, 554. 10.1186/s13059-014-0554-4.

122. Suzuki, K., Bose, P., Leong-Quong, R.Y., Fujita, D.J., and Riabowol, K. (2010). REAP: A two minute cell fractionation method. BMC Res Notes 3, 294. 10.1186/1756-0500-3-294.

123. Artan, M., Hartl, M., Chen, W., and de Bono, M. (2022). Depletion of endogenously biotinylated carboxylases enhances the sensitivity of TurboID-mediated proximity labeling in Caenorhabditis elegans. J Biol Chem 298, 102343. 10.1016/j.jbc.2022.102343.

124. Rappsilber, J., Mann, M., and Ishihama, Y. (2007). Protocol for micro-purification, enrichment, pre-fractionation and storage of peptides for proteomics using StageTips. Nat Protoc 2, 1896–1906. 10.1038/nprot.2007.261.

125. Tyanova, S., Temu, T., and Cox, J. (2016). The MaxQuant computational platform for mass spectrometry-based shotgun proteomics. Nat Protoc 11, 2301–2319. 10.1038/nprot.2016.136.

126. UniProt https://www.uniprot.org/.

127. R: The R Project for Statistical Computing https://www.r-project.org/.

128. Madern, M. (2023). Cassiopeia_LFQ.

129. Chen, E.Y., Tan, C.M., Kou, Y., Duan, Q., Wang, Z., Meirelles, G.V., Clark, N.R., and Ma’ayan, A. (2013). Enrichr: interactive and collaborative HTML5 gene list enrichment analysis tool. BMC Bioinformatics 14, 128. 10.1186/1471-2105-14-128.

130. Kuleshov, M.V., Jones, M.R., Rouillard, A.D., Fernandez, N.F., Duan, Q., Wang, Z., Koplev, S., Jenkins, S.L., Jagodnik, K.M., Lachmann, A., et al. (2016). Enrichr: a comprehensive gene set enrichment analysis web server 2016 update. Nucleic Acids Res 44, W90–97. 10.1093/nar/gkw377.

131. Xie, Z., Bailey, A., Kuleshov, M.V., Clarke, D.J.B., Evangelista, J.E., Jenkins, S.L., Lachmann, A., Wojciechowicz, M.L., Kropiwnicki, E., Jagodnik, K.M., et al. (2021). Gene Set Knowledge Discovery with Enrichr. Curr Protoc 1, e90. 10.1002/cpz1.90.

132. Perez-Riverol, Y., Csordas, A., Bai, J., Bernal-Llinares, M., Hewapathirana, S., Kundu, D.J., Inuganti, A., Griss, J., Mayer, G., Eisenacher, M., et al. (2019). The PRIDE database and related tools and resources in 2019: improving support for quantification data. Nucleic Acids Res 47, D442–D450. 10.1093/nar/gky1106.

133. Neuhold, J., Radakovics, K., Lehner, A., Weissmann, F., Garcia, M.Q., Romero, M.C., Berrow, N.S., and Stolt-Bergner, P. (2020). GoldenBac: a simple, highly efficient, and widely applicable system for construction of multi-gene expression vectors for use with the baculovirus expression vector system. BMC Biotechnology 20, 26. 10.1186/s12896-020-00616-z.

134. Ehrmann, J.F., Grabarczyk, D.B., Heinke, M., Deszcz, L., Kurzbauer, R., Hudecz, O., Shulkina, A., Gogova, R., Meinhart, A., Versteeg, G.A., et al. (2023). Structural basis for regulation of apoptosis and autophagy by the BIRC6/SMAC complex. Science 379, 1117–1123. 10.1126/science.ade8873.

135. GATK https://gatk.broadinstitute.org/hc/en-us.

136. Benjamin, D., Sato, T., Cibulskis, K., Getz, G., Stewart, C., and Lichtenstein, L. (2019). Calling Somatic SNVs and Indels with Mutect2. Preprint at bioRxiv, 10.1101/861054 10.1101/861054.

137. MutationalPatterns (2024). (UMCU Genetics).

138. Rozen, S.G. (2024). mSigAct.

139. DCC Data Releases | ICGC Data Portal https://dcc.icgc.org/releases/PCAWG.

140. The Cancer Genome Atlas Program (TCGA) - NCI (2022). https://www.cancer.gov/ccg/research/genome-sequencing/tcga.

